# Adolescent prodromal phenotypes of psychiatric disorders: insights from a novel DISC1-perturbed monkey model

**DOI:** 10.64898/2026.10.11.758339

**Authors:** Ziguo Lan, Hirotaka Onoe, Tomoyuki Tsukiyama, Chizuru Iwatani, Chih-Yang Chen, Keisuke Okita, Koko Ishizuka, Ryo Endo, Risako Nakai, Keiko Imamura, Keigo Hikishima, Hideaki Tsuchiya, Masataka Nakaya, Yang Fang, Atsushi Kobayashi, Yuta Shinto, Reona Yamaguchi, Mami Matsunaga, Kengo Oe, Koichi Omori, Yuji Ozeki, Toshifumi Tomoda, Kenichiro Miura, Shin-ichi Urayama, Naoya Oishi, Tomohisa Okada, Motomasa Tanaka, Haruhisa Inoue, Jun Takahashi, Akira Sawa, Tadashi Isa

**Author notes:** Equally contributed (Ziguo Lan, Hirotaka Onoe, Tomoyuki Tsukiyama).

## Abstract

Neurodevelopmental deficits frequently lead to mild but significant adolescent social and cognitive changes, representing prodromal manifestations of adult-onset psychiatric disorders in a cross-disease manner. Although experimental medicine utilizes animal models to address the mechanisms, rodent brains lack critical structures for human higher brain functions such as the dorsolateral prefrontal cortex. To fill this translational gap, we generated monkeys with an impaired *DISC1* gene locus, targeting the multifunctional protein essential for neurodevelopment and involved in transdiagnostic mental health risks. Here we analyze the F0 animals by minimizing mosaicism. At ages equivalent to human adolescence, these monkeys exhibited blunted responsiveness to social stimuli and increased internal drives, mirroring altered social behaviors in the prodromal stage in humans. Resting-state functional MRI data indicated disrupted global functional connectivity and reorganization of network topology, marked by a shift in the dominance of brain hubs from prefrontal regions to sensory areas. This primate model provides a unique platform for studying psychiatric disorders, such as schizophrenia, across the developmental trajectory.

## INTRODUCTION

Unlike cancers or metabolic disorders, modeling neuropsychiatric conditions in animals faces fundamental challenges. First, the dorsolateral prefrontal cortex (DLPFC), critical for schizophrenia (SZ) (*1*, *2*), is absent in rodents (*3*). However, key brain regions implicated in these disorders are uniquely developed in primates, highlighting the importance of non-human primate models. Second, many adult-onset psychiatric disorders arise from neurodevelopmental disturbances (*4*). Therefore, understanding the pathobiology prior to onset is crucial for effective treatment, including prophylactic interventions (*5*, *6*).

Accordingly, it is essential to study prodromal stages to elucidate early pathobiology and enable preventive interventions. Third, current diagnostic criteria prioritize reliability over biological validity (*7–10*), meaning that animal models based solely on genes identified through genome-wide association studies (GWASs), which rely on such criteria (e.g., DSM), should be interpreted with caution.

DISC1, an intracellular hub protein, has been widely studied due to its role in cognitive and emotional regulation (*11*, *12*). Its disruption was first identified in a Scottish pedigree with a high incidence of adult-onset severe mental illnesses, including SZ, bipolar disorder, and major depressive disorder (*13*, *14*). Although its direct genetic link to SZ remains debated, DISC1 interacts with numerous proteins associated with psychiatric risk (*15–17*), supporting its importance in higher brain functions. Mouse models with DISC1 loss-of-function have demonstrated that disruptions during neurodevelopment can lead to adult behavioral abnormalities relevant to psychiatric conditions (*18–21*).

To fill the translational gap between rodents and humans, we generated cynomolgus monkeys (*Macaca fascicularis*) with an impairment in the *DISC1* gene locus (DISC1-LI monkey), which may help elucidate the pathophysiology and endophenotypes of neuropsychiatric disorders. To efficiently generate DISC1-LI monkeys suitable for F0 phenotypic evaluation, we applied the optimized genome-editing strategy to suppress mosaic mutations (*22*). We targeted critical developmental windows relevant to the prodromal stages of adult-onset psychiatric disorders. We observed behavioral and neurocircuitry alterations which mirror prodromal manifestations, providing a unique platform for studying psychiatric disorders, such as SZ, across the developmental trajectory.

## RESULTS

### Generation of DISC1-LI monkeys

We generated monkeys with an impaired *DISC1* gene locus by co-injecting a high concentration of gRNA/Cas9 protein RNP complexes with sperm into oocytes during the metaphase II-stage, minimizing mosaicism and enabling analysis of F0 animals (*22*). Among 9 embryos transferred into surrogate mothers for the Exon 2-5 deletion group, 2 embryos successfully implanted, while in the Exon 2-Intron 8 deletion group, 3 of 7 embryos were successfully implanted. Consequently, a total of 5 fetuses were delivered (**Fig. 1, A to C**).

**Fig. 1.**
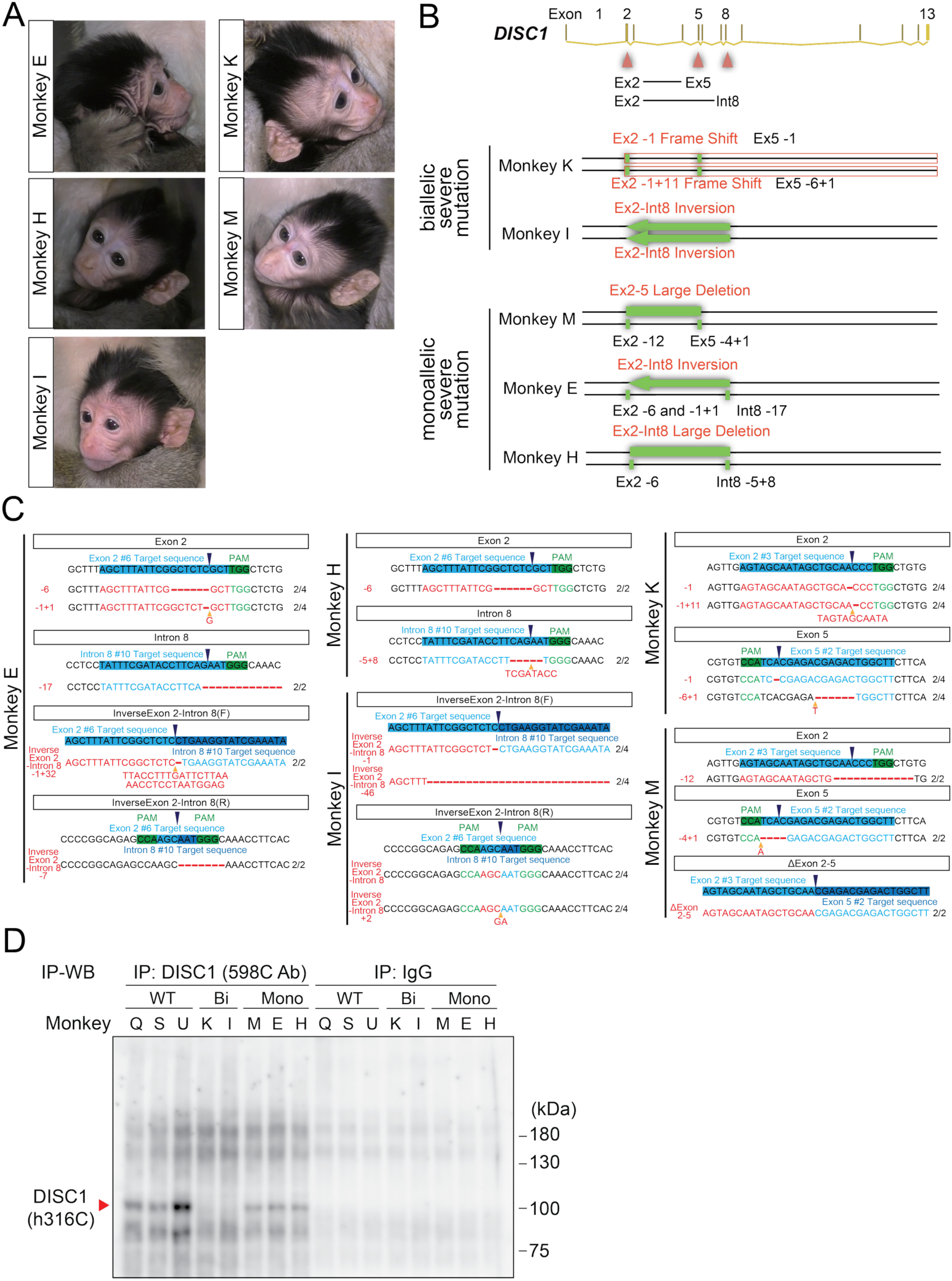
Genotyping DISC1-LI monkeys. (**A**) Five F0 DISC1-LI monkeys. (**B**) The *DISC1* gene in each monkey. (**C**) Sequences of sgRNA targets in the genomes of the DISC1-LI monkeys. **(D)** Immunoprecipitation combined with western blotting (IP-WB) for DISC1 in iPSCs derived from wild type (WT) and DISC1-LI monkeys. Full-length DISC1 (∼100 kDa) was detected in the cells of WT but not in biallelic mutation subjects (Monkey K and I). In the cells from monoallelic mutation subjects (Monkey M, E and H), a full-length DISC1 signal remained detectable at approximately 50% of the WT level. IgG was used as a negative control.

Genotyping through DNA sequencing revealed that Monkey K had frameshift mutations in both alleles, whereas Monkey I possessed an Exon 2-Intron 8 inversion in both alleles. In these two fetuses, both alleles (DISC1*^Bi^*) were severely disrupted and likely resemble complete DISC1-perturbed monkeys (**Fig. 1 and fig. S4**). In contrast, Monkey M had a large deletion of Exon 2-5 only in one allele and a missense mutation in the other; Monkey E exhibited an Exon 2-Intron 8 inversion in one allele and two missense mutations in the other; and Monkey H harbored a large deletion of Exon 2-Intron 8 in one allele and a missense mutation in the other. Thus, these three fetuses may harbor a severe disruption only in a single allele (DISC1*^Mono^*) (**Fig. 1 and fig. S4**).

The depletion of DISC1 protein was validated in these DISC1-LI monkeys. According to the genomic data, we predicted that Monkeys K and I would entirely lack DISC1 protein, whereas Monkeys M, E and H might maintain approximately half the level of DISC1 protein. To analyze DISC1 protein levels, we generated induced pluripotent stem cells (iPSCs) from each individual and examined the expression of endogenous DISC1 (**Fig. 1D and fig. S5**). In cells from two subjects (Monkey K and I), full-length DISC1 protein was completely undetectable. In cells from the other three subjects (Monkey M, E and H), DISC1 protein of nearly full-length size remained detectable but was reduced to approximately half the level detected in WT monkeys. These data are consistent with predicted outcomes from the genomic manipulation.

### Altered socio-emotional responses to social stimuli in DISC1-LI monkeys

Mild but significant adolescent socio-emotional alterations are frequently observed in the prodromal stages of adult-onset psychiatric disorders, such as SZ. To examine whether DISC1-LI monkeys display these alterations in adolescence, we conducted the human intruder test (HIT), a structured paradigm assessing responses to social stimuli (**Fig. 2A**). During the Alone phase, external behaviors were minimal in both genotypes, establishing a comparable baseline in the test environment. Upon social challenge in the Profile and Stare phases, however, DISC1-LI monkeys displayed markedly attenuated external responses relative to WT controls (**Fig. 2B**), suggesting diminished reactivity to the intruder’s presence. Conversely, these monkeys exhibited significantly prolonged internal behavior across all experimental phases (**Fig. 2C**). These patterns were further corroborated by standardized behavioral scores (**Fig. 2, D and E**).

**Fig. 2.**
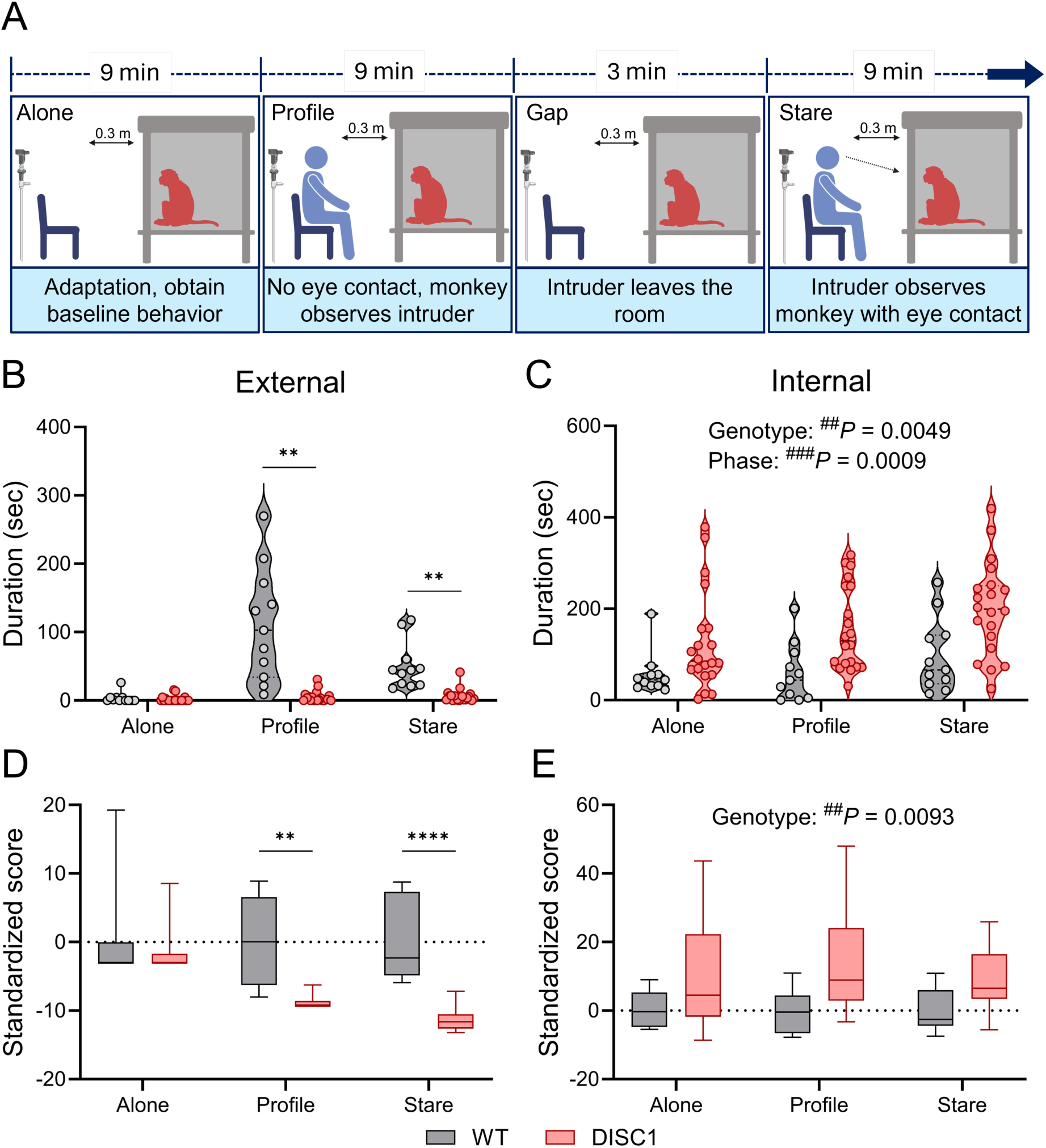
DISC1-LImonkeys showed social deficits in the human intruder test. (**A**) Test protocol (created with BioRender.com). WT, *N* = 4 (total trials = 11); DISC1-LI, *N* = 5 (total trials = 20). (**B**) Duration of external behavior. DISC1-LI monkeys showed shorter external responses during *Profile* (\*\**P* = 0.0017) and *Stare* (\*\**P* = 0.0017) phases compared to WT (Genotype × Phase, ^####^*P* < 0.0001; Genotype, ^####^*P* < 0.0001; Phase, ^####^*P* < 0.0001). (**C**) Duration of internal behavior. DISC1-LI monkeys exhibited prolonged internal behavior across all phases compared to WT monkeys (Genotype, ^##^*P* = 0.0049; Phase, ^###^*P* = 0.0009). (**D**) Standardized score of external behavior. DISC1-LI monkeys show significantly lower scores during *Profile* (\*\**P* = 0.0011) and *Stare* (\*\*\*\**P* < 0.0001) phases (Genotype × Phase, ^####^*P* < 0.0001; Genotype, ^####^*P* < 0.0001; Phase, ^###^*P* = 0.0001). (**E**) Standardized score of internal behavior. DISC1-LI monkeys show significantly higher scores (Genotype: ^##^*P* = 0.0093). Two-way RM ANOVA with Greenhouse-Geisser correction, followed by Tukey’s HSD test.

A closer examination of specific behavioral categories further clarified these disparities. Among external behaviors, anxiety-related actions were significantly reduced in DISC1-LI monkeys during the *Profile* phase (**Fig. S6A**), while excitation and defense behaviors were markedly higher in WT monkeys during both *Profile* and *Stare* phases (**Fig. S6, B and C**). Submission behaviors showed no significant intergroup differences (**Fig. S6D**), whereas the frequency of affiliative behavior was decreased in DISC1-LI monkeys during both *Profile* and *Stare* phases (**Fig. S6E**). Standardized scores across all external behavior categories further supported these findings (**Fig. S7**). For internal behaviors, although stereotyped behavior did not reach statistical significance (**Fig. S6F**), DISC1-LI monkeys exhibited elevated exploratory behavior in both frequency and duration, along with increased frequency of other behaviors, particularly in the *Profile* phase (**Fig. S6, G and H**).

We next asked whether the extent of the perturbation of DISC1 protein influenced these behavioral phenotypes by comparing HIT performance between monoallelic (DISC1*^Mono^*, Monkey M, E and H) and biallelic (DISC1*^Bi^*, Monkey K and I) mutation individuals. The DISC1*^Mono^* and DISC1*^Bi^* groups exhibited similar responses to intruder stimuli (**Fig. S8, A to E**) and comparable internal behaviors (**Fig. S8, F to H**).

### Alterations in functional brain connectivity and network architecture in DISC1-LI monkeys

We hypothesized that the aberrant social behaviors might reflect altered functional brain connectivity and network architecture elicited by DISC1 perturbation. Accordingly, we analyzed functional connectivity (FC) derived from resting-state functional MRI (rsfMRI) data. DISC1-LI monkeys exhibited significantly decreased overall connectivity strength and density compared to the WT group, particularly in intra-regional frontal connections, as well as inter-regional connections between the frontal lobes and motor or temporal areas. These affected regions included the DLPFC and posterior cingulate cortex (PCC), suggesting region-specific vulnerability (**Fig. 3, A and B**).

**Fig. 3.**
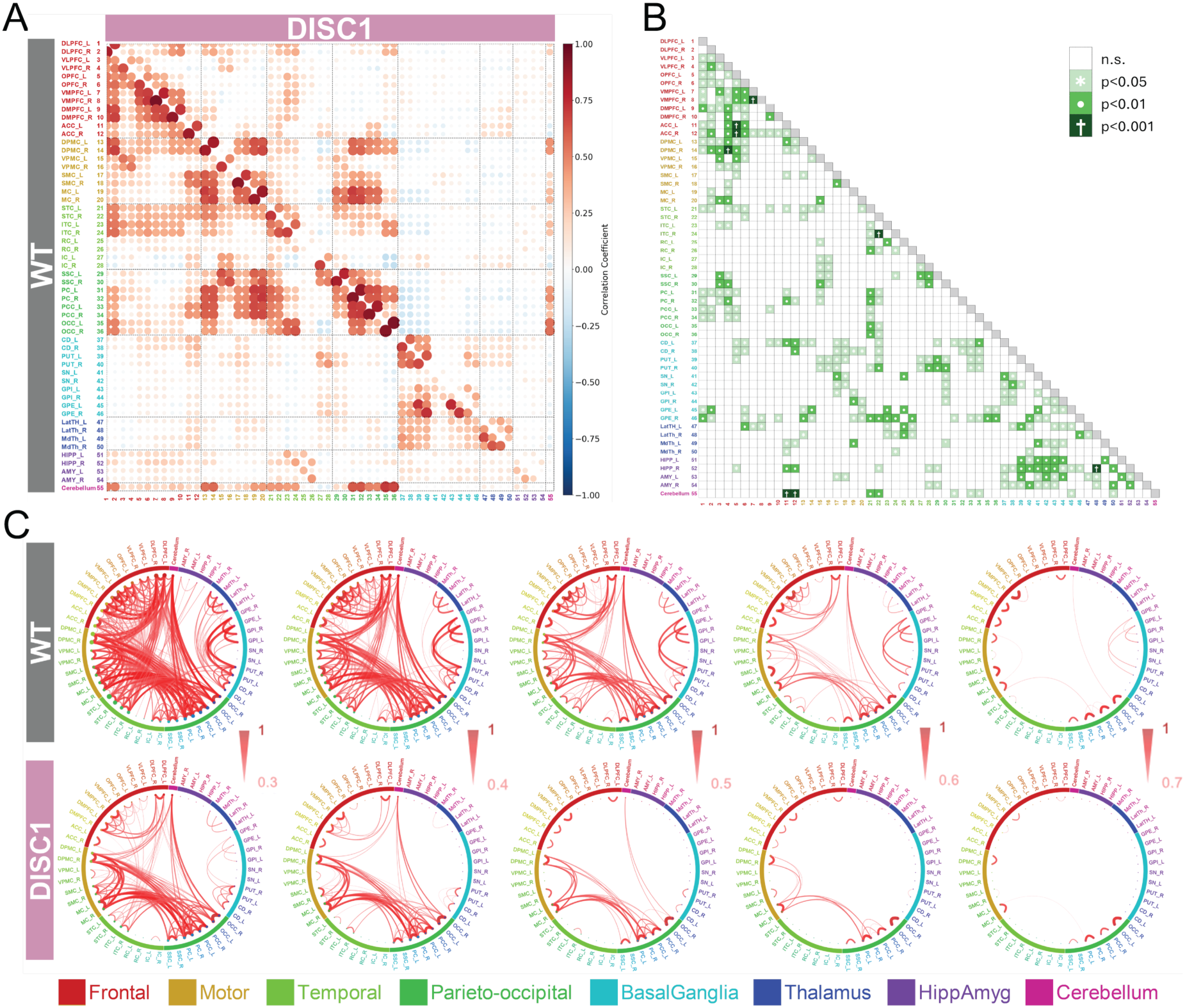
Global functional connectivity (FC) in DISC1-LI monkeys decreased. (**A**) FC matrices for WT (*N* = 6) and DISC1-LI (*N* = 5) monkeys. (**B**) Statistical comparison of FC z-score between WT and DISC1-LI groups. t-test with Welch’s correction. (**C**) Chord diagrams of FC at thresholds of 0.3 to 0.7.

We further examined the network architecture across a range of correlation thresholds to evaluate topological robustness (**Fig. 3C**). At lower thresholds, WT controls displayed densely interconnected networks with profuse intra- and inter-regional connectivity, whereas DISC1-LI monkeys showed a markedly more fragmented topology with diminished interhemispheric and long-range connections. As the threshold increased, progressive network sparsification occurred in both groups, but this decline was considerably more pronounced in DISC1-LI monkeys. At higher thresholds, only minimal, predominantly localized connectivity was preserved in DISC1-LI monkeys, in sharp contrast to the relatively intact networks maintained in WT controls (**Fig. 3C**).

To exclude the possibility that these connectivity changes might be elicited by specific anesthetic agents, we compared the FC under alfaxalone and ketamine anesthesia. The FC patterns for both WT (**Fig. S9A**) and DISC1-LI monkeys (**Fig. S9B)** were equivalent between these two anesthetic conditions, indicating that the observed differences were attributable to DISC1 perturbation.

We also asked whether the degree of perturbation in DISC1 protein influenced these connectivity changes by comparing individuals with monoallelic perturbation (DISC1*^Mono^*, Monkey M, E and H) with those with biallelic perturbation (DISC1*^Bi^*, Monkey K and I). Both groups displayed a robust reduction in the functional connectivity compared to WT monkeys, although the severity of the alterations differed (**Fig. S10**). DISC1-LI monkeys with monoallelic perturbation exhibited more pronounced reductions in long-range connections between frontal-motor and frontal-temporal regions (**Fig. S10A**), whereas DISC1-LI monkeys with biallelic perturbation showed decreased FC in key frontal-related circuits, while the overall network integrity appeared paradoxically less compromised (**Fig. S10B**).

### Alterations in local network efficiency and small-world organization in DISC1-LI monkeys

Having established broad FC alterations, we further determined whether global and local network organizations were also disrupted in DISC1-LI monkeys using small-world network analysis. While the clustering coefficient (*C_p_*) did not differ between groups (**Fig. S11A**), DISC1-LI monkeys showed significantly shorter characteristic path lengths (*L_p_*) at low sparsity thresholds (**Fig. S11B**). In DISC1-LI monkeys, the normalized clustering coefficient (*γ*) was reduced (**Fig. S11C**), while normalized path lengths (*λ*) showed no significant difference (**Fig. S11D**), resulting in consistently lower small-worldness (*σ*) compared to the WT group (**Fig. S11E**). Furthermore, even though global efficiency (*E_glob_*) was comparable between the two groups (**Fig. S11F**), local efficiency (*E_loc_*) was significantly diminished in the DISC1 group at low sparsity thresholds (**Fig. S11G**), highlighting the potential impairment of local information transfer. Together, these results suggest a disrupted balance between local clustering and global integration. Although an area under the curve (AUC) analysis did not reach statistical significance between the two groups (**Fig. S11H**), the overall pattern points to a complex reorganization of brain networks in DISC1-LI monkeys.

### Fewer prefrontal and more dominant sensory hubs: an altered network centrality landscape in DISC1-LI monkeys

The FC analysis revealed widespread connectivity alterations, while the small-world analysis further indicated altered local network organization. To identify which specific regions drove these alterations, we assessed eigenvector centrality at FC sparsity thresholds of 0.05, 0.1, 0.2, and 0.3. At the lowest sparsity threshold, both groups commonly exhibited prominent hub activity in motor (primary motor cortex, MC; dorsal premotor cortex, DPMC) and parietal regions (parietal cortex, PC; PCC). Whereas in WT monkeys the DLPFC displayed comparatively high eigenvector centrality at this level, emerging as a significant hub within the frontal lobe (**Fig. 4A**), and robustly maintaining this role as network density increased (**Fig. 4, A to D**). In contrast, DISC1-LI monkeys displayed a dramatically altered network hub configuration. The DLPFC of these monkeys exhibited diminished eigenvector centrality with limited connectivity across all network densities explored (**Fig. 4, E to H**), signifying a profound loss of prefrontal dominance in the global network. This deficit was accompanied by a striking redistribution of network influence toward sensory-related regions, where the somatosensory cortex (SSC) and occipital cortex (OCC) specifically demonstrated consistently elevated centrality, functioning as dominant hubs within the DISC1-LI monkeys’ network (**Fig. 4, E to H**). Such reorganization indicates a redistribution of network influence away from prefrontal regions toward sensory-related areas, reflecting a disrupted balance in cortical hub architecture. Given the established role of the DLPFC in higher-order cognition and social behavior, these network-level alterations may be associated with the socio-emotional behavioral phenotypes observed in these animals (**Fig. 2**).

**Fig. 4.**
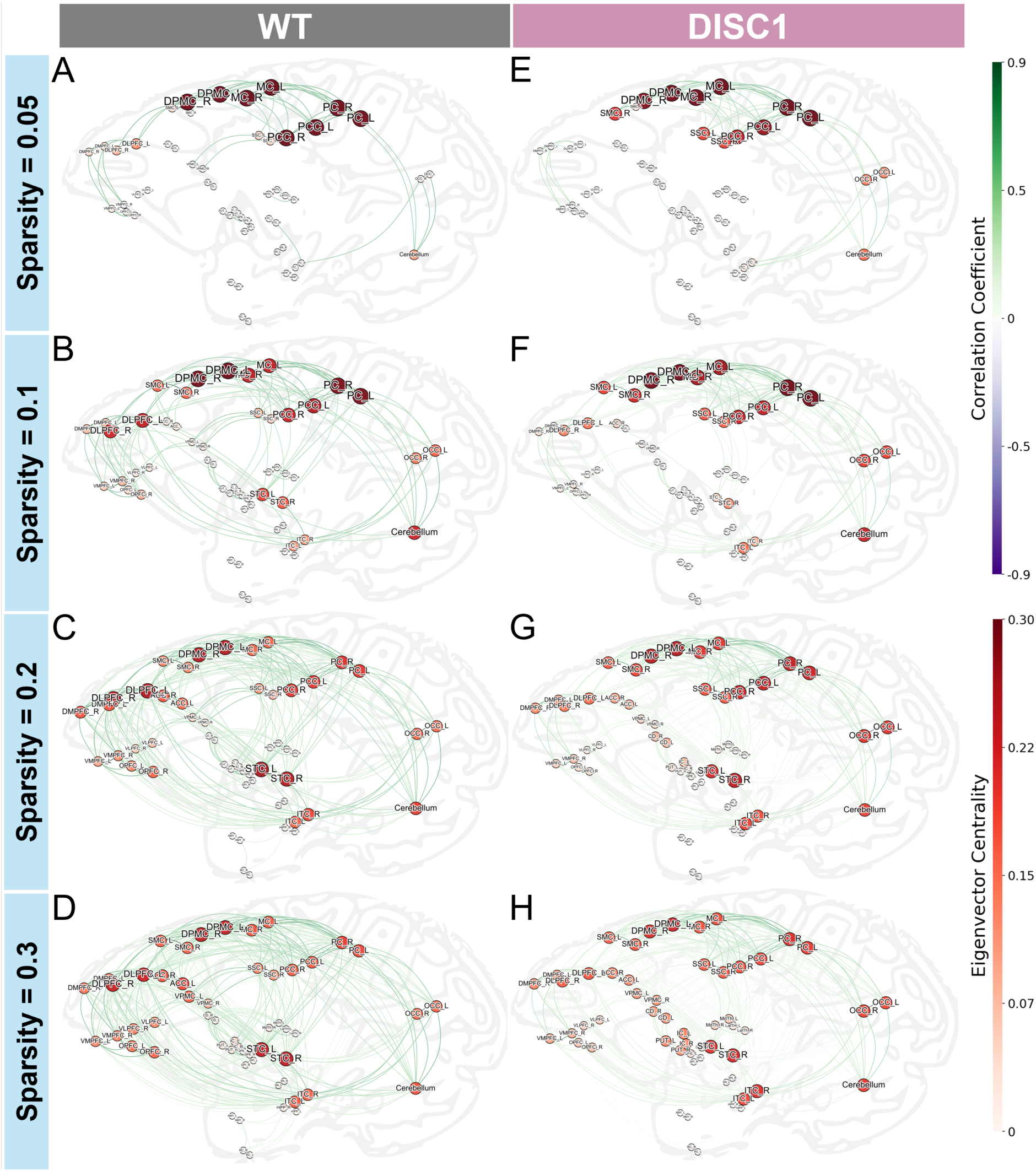
Eigenvector centrality architecture in the brain network of DISC1-LI monkeys was altered. The brain network at sparsity thresholds of 0.05, 0.1, 0.2, and 0.3 for WT (*N* = 6) (**A** to **D**), and DISC1-LI (*N* = 5) (**E** to **H**) monkeys. Node size and color indicate eigenvector centrality, with larger and redder nodes signifying higher centrality values. Edges represent functional connections, with edge color intensity corresponding to correlation strength.

## DISCUSSION

In this study, we established a genetically engineered monkey model with targeted disruptions in the gene encoding DISC1, a critical molecular hub involved in neurodevelopmental processes (*23–26*). To minimize mosaicism, we generated F0 animals by co-injecting gRNA/Cas9 protein RNP complexes along with sperm into cynomolgus metaphase II-stage oocytes. DISC1-LI monkeys displayed increased internal self-directed behaviors in adolescence, coinciding with alterations in neural connectivity, particularly reduced prefrontal connectivity.

Monkey models for neuropsychiatric conditions have recently been generated to address the pathobiological mechanisms of child-onset developmental disorders and geriatric neurodegenerative diseases by manipulating causal genes for specific diseases (*27–39*). In addition, pharmacological administrations, such as the use of ketamine, have been used to mimic the symptomatic changes of adult-onset neuropsychiatric conditions (*40–45*). In contrast, we targeted a lead molecule that can drive adult-onset neuropsychiatric manifestations and neurophysiological changes relevant to SZ (e.g., event-related potential P300) to establish a novel primate model (*14*, *46*). Perturbation of DISC1 in mice has also been known to result in late-onset behavioral changes after puberty, mirroring human adult-onset psychiatric disorders (*11*, *16*, *18*). Our present data examining adolescent brain and behavioral changes elicited by DISC1 loss-of-function provide novel insights that were not addressed by previous primate models for neuropsychiatric conditions.

Our findings in DISC1-LI monkeys at ages corresponding to human adolescence (approximately 8-15 years old) reveal a parallel to those reported in adolescent longitudinal human studies. The Philadelphia Neurodevelopmental Cohort (PNC) demonstrates that psychosis spectrum symptoms can emerge in youth and often persist or worsen over time, with early cognitive and social deficits predicting a trajectory towards severe psychotic disorders, such as SZ (*47–49*). The cohort study also reports that at-risk subjects or those in the prodromal stages exhibit aberrant functional connectivity involving the frontal regions (*50*). In this context, the phenotypes observed in adolescent DISC1-LI monkeys resemble those in the prodromal stages of adult-onset psychiatric disorders such as SZ.

Based on DISC1 protein levels, the five DISC1-LI monkeys can be divided into two subgroups: Monkey K and I (biallelic, complete protein deletion) and Monkey M, E, and H (monoallelic, partial protein deletion). However, within the analyses conducted thus far, we did not observe robust phenotypic differences between these two subgroups. Due to the missense mutations in the remaining *DISC1* allele in the latter group, both subgroups may be functionally equivalent to a complete loss-of-function state. Alternatively, a partial DISC1 loss-of-function may be sufficient to elicit the observed phenotypes within our assays. While this uncertainty represents a minor limitation of the present modeling, future molecular dissection of all five subjects may provide deeper mechanistic insights.

Although the present study focuses on a cross-sectional, adolescent assessment of these DISC1-LI monkeys, longitudinal tracking of these subjects will be essential in future research. Given that these subjects display neurocircuitry and behavioral alterations that are reminiscent of those found in the prodromal stages of adult-onset psychiatric disorders, such as SZ, a longitudinal assessment in the coming 2-4 years may provide us with an opportunity to observe the conversion from prodromal stages to the full development of a disorder in young adulthood. An expansion of phenotypic outcome measures, including eye movements, sleep architecture, prepulse inhibition (PPI), mismatch negativity (MMN), event-related potentials N50 and P300, decision-making, attentional control, self-identity, and proxies for hallucinations and delusions will further enrich the translational utility of this model.

Furthermore, comparing published brain imaging data from the Scottish pedigree with the phenotypes from these DISC1-LI monkeys later in adult stages will be of great importance (*51–54*). Recently, Zhou et al. (*55*) reported the behavioral and physiological phenotypes of 3 adult F1 cynomolgus monkeys with disruption of exon 9 of the *DISC1* gene. Further in-depth analysis and follow-up of these non-human primate resources will shed light on how adult-onset psychiatric disorders, such as SZ, emerge from the developmental process through the prodromal stages.

In the long term, elucidating the DISC1 protein interactome and examining how protein-protein interactions and downstream signaling pathways are disrupted could provide molecular insights into observed neural and behavioral phenotypes shared between this primate model and patients. Finally, experimental therapeutic strategies that target network dysfunction, such as deep brain stimulation (DBS) of the substantia nigra pars reticulata (SNr), are currently underway for SZ patients (*56*, *57*). Optogenetics and chemogenetic circuit interventions in DISC1-LI monkeys and clinical DBS will be mutually informative, opening a new chapter of translational approaches for neuropsychiatric disorders.

## MATERIALS AND METHODS

### Study approval

We followed the Reporting in Vivo Experiments (ARRIVE) guidelines developed by the National Centre for the Replacement, Refinement & Reduction of Animals in Research (NC3Rs). We also followed “The Act on Welfare and Management of Animals” from Ministry of the Environment, “Fundamental Guidelines for Proper Conduct of Animal Experiment and Related Activities in Academic Research Institutions” under the jurisdiction of the Ministry of Education, Culture, Sports, Science and Technology, and “Guidelines for Proper Conduct of Animal Experiments” from Science Council of Japan. All animal experimental procedures were approved by the Committee of Animal Experimentation (approval number: Med Kyo19599, 20035, 21013, 22030, 23024, 24055, 25518 and 26066) and the Recombinant DNA Technology Committee (approval number: 200429 and 250421) of Kyoto University, and Animal Care and Use Committee (approval number: 2019-10-4, 2022-11-4 and 2024-9-10) and the Recombinant DNA Experiment Safety Committee (approval number: 31-30 and 2024-026) of Shiga University of Medical Science.

### Animals

For oocyte collection, female cynomolgus monkeys (*Macaca fascicularis*), aged 4 to 13 years, were selected for this study. The animals were maintained under a 12-hour light/dark cycle, with artificial lighting provided from 8:00 AM to 8:00 PM. Each monkey was provided with 20 g/kg of body weight of commercial pellet feed (CMK-1; CLEA Japan) in the morning, supplemented with 20–50 g of sweet potato in the afternoon. Water was available *ad libitum*. Temperature and humidity in the animal rooms were maintained at 25 ± 2°C and 50 ± 5%, respectively.

### Generation of DISC1-LI monkeys

#### gRNA design and selection

To efficiently generate DISC1-LI monkeys with extensive mutations, guide RNAs (gRNAs) were designed targeting Exon 2 and Exon 5, or Exon 2 and Intron 8, regions where highly specific gRNAs could be identified. Six candidate gRNAs were initially designed for Exon 2, four for Exon 5, and five for Intron 8. However, one gRNA targeting Exon 2 and two targeting Exon 5 were excluded from the candidates due to the presence of polymorphisms in their sequences, as identified by DNA sequence analysis in WT monkeys (**Fig. S1**). From the remaining five candidate gRNAs in Exon 2, two in Exon 5, and five in Intron 8, pairs were selected using a single-strand annealing (SSA) assay (*58*).

#### Single-strand annealing (SSA) assay

pX330 was obtained from Addgene (Plasmid #42230) (*59*). To construct pX330-DISC1, the oligonucleotides listed in Supplementary Table 1 were annealed and ligated into the BbsI site of pX330. To construct pCAG-EGxxFP-DISC1, an amplified PCR product was cloned into the EcoRI-NheI sites of pCAG-EGxxFP (*58*), using the primers listed in Supplementary Table 1. 293FT cells were co-transfected with pX330-DISC1 and pCAG-EGxxFP-DISC1.

Two days after incubation in Dulbecco’s modified Eagle’s medium (Sigma) supplemented with 10% fetal bovine serum (Sigma), the cells were harvested and analyzed by flow cytometry. As a criterion for gRNA selection, priority was assigned to those targeting Exon 2 for maximal functional disruption, with favorable results demonstrated in the SSA assay. As a result, the following pairs were identified: Exon 2 #3 with Exon 5 #2, and Exon 2 #6 with Intron 8 #10 (**Fig. S2**).

#### Oocyte and sperm collection

Two weeks after the subcutaneous injection of 0.9 mg of a gonadotropin-releasing hormone antagonist (Leuplin for Injection Kit, Takeda Chemical Industries), a micro infusion pump (iPRECIO SMP-200, ALZET Osmotic Pumps) with 15 IU/kg human follicle-stimulating hormone (hFSH, Gonapure Injection, Asuka Pharmaceutical) was surgically implanted subcutaneously under anesthesia. The pump administered 7 µl/h for 10 days. After the hFSH treatment, 400 IU/kg human chorionic gonadotropin (hCG, Gonatropin, Asuka Pharmaceutical) was injected intramuscularly. Forty hours after the hCG treatment, oocytes were collected by follicular aspiration using a laparoscope (LA-6500, Machida Endoscope). Fresh sperm were collected by electric stimulation of the penis without anesthesia.

#### Intracytoplasmic sperm injection (ICSI)

Cumulus-oocyte complexes (COCs) were recovered in alpha modification of Eagle’s medium (MP Biomedicals, Solon), containing 10% serum substitute supplement (Irvine Scientific).

Cumulus cells were enzymatically removed from the COCs by treatment with 0.5 mg/ml hyaluronidase (Sigma Chemical). ICSI was carried out on metaphase II (MII)-stage oocytes in mTALP containing HEPES with a micromanipulator. Following ICSI, embryos were cultured in CMRL Medium-1066 (Invitrogen) supplemented with 20% bovine serum (Invitrogen) at 38°C in 5% CO_2_ and 5% O_2_. To construct RNP complexes, the crRNA, tracrRNA, and HiFi Cas9 proteins were purchased from Integrated DNA Technologies, and the gRNA/Cas9 protein RNP complexes were assembled according to the manufacturer’s instructions.

To confirm the cleavage efficiency of selected gRNA pairs in fertilized monkey eggs, gRNA/Cas9 protein RNP complexes along with sperm were co-injected into cynomolgus MII-stage oocytes. A high concentration (200 ng/µl) of Cas9 protein was employed to ensure complete mutation induction in single-cell embryos, thereby minimizing the risk of mosaicism to enable analysis in F0 animals (*22*). Furthermore, to achieve effective large-scale deletions between gRNA pairs, the injection solutions were supplemented with 50 ng/µl of repair templates. These repair templates included genomic sequences of 80 base pairs in total, with 40 base pairs flanking both the upstream and downstream regions of the indel induction sites. MII oocytes and sperm were washed in a drop of 200 ng/µl RNP complex supplemented with 50 ng/µl repair templates in Opti-MEM (Invitrogen), and ICSI was performed in the drop.

#### Embryo culture and validation of mutation induction

For the Exon 2-5 deletion experiment, a total of 38 oocytes were injected, of which 36 survived and 17 developed to the blastocyst stage. In the Exon 2-Intron 8 deletion experiment, 29 oocytes were injected, 23 survived, and 13 reached the blastocyst stage. During embryonic development, three embryos at the 7-9 cell stage were sampled from each group, and individual blastomeres were genotyped. DNA sequencing analysis identified indel mutations, large deletions, or inversions near the Cas9 cleavage site in all embryos, demonstrating the high efficacy of the gRNAs (**Fig. S3**).

#### Embryo transfer

When the embryos developed into expanded blastocysts, a single embryo was transferred into each appropriate recipient female (*60*, *61*). Embryos were aspirated into a catheter (ETC3040SM5-17; Kitazato Medical Service) under a stereomicroscope. The catheter was inserted into the oviduct of the recipient via the fimbria under laparoscopic guidance, and the cultured embryo was transferred along with a minimal volume of medium. Pregnancy was determined by ultrasonography 30 days following ICSI. To determine the genotypes of the offspring, genomic DNA was extracted from blood samples.

### iPSCs generation

iPSCs were generated from 0.5–1.5 mL of peripheral blood. Peripheral blood mononuclear cells were isolated by density gradient centrifugation using Percoll (Merck) and transfected with a combination of plasmid vectors (OCT3/4, SOX2, KLF4, L-MYC, LIN28, EBNA1 and a dominant negative form of mTrp53) as described previously (*62*). Up to 10 clones per monkey were selected for further experiments.

### Validation of loss-of DISC1 protein

Cynomolgus monkey iPSCs at day 0 of differentiations were lysed in immunoprecipitation buffer (10 mM HEPES pH7.4, 150 mM NaCl, 1% NP-40, 1 mM EDTA, 1 mM DTT) supplemented with complete EDTA-free protease inhibitor cocktail (Nacalai tesque). Dynabeads Protein G magnetic beads were incubated with anti DISC1 antibody (h598C) or normal rabbit IgG as a negative control and cross-linked with 25 mM dimethyl pimelimidate (DMP) (Thermofisher) in 0.2 M triethanolamine (pH 8.2) for 60 min at room temperature (*63*). Excess DMP was quenched with 0.2 M ethanolamine in phosphate buffered saline and beads were washed with wash buffer (50 mM Tris-Cl pH7.5, 150 mM NaCl, 0.5% NP-40), followed by incubation with cell lysate for overnight at 4°C. The immunoprecipitates were subjected to western blotting using anti DISC1 antibody (h316C) (*63*).

### Behavioral test

#### Human intruder test

9 cynomolgus monkeys including 4 WT (3-4.5 years old) and 5 DISC1-LI individuals (3-3.5 years old), were assessed using the human intruder test (HIT). All subjects were housed individually in a light- and temperature-controlled colony room with feeding as described above and *ad libitum* access to water. None of the subjects had any prior implantation or cranial surgery.

The experimental paradigm consisted of four phases: *Alone*, *Profile*, *Gap*, and *Stare*, each performed sequentially in a controlled environment (**Fig. 2A**). During the initial *Alone* phase (9 minutes), the subject adapted to the environment, and baseline behavior was recorded. In the *Profile* phase (9 minutes), the intruder sat 0.3 m away, avoiding direct eye contact by looking somewhere away from monkey’s head. After a brief *Gap* phase (3 minutes), where the intruder left the room, the *Stare* phase (9 minutes) involved the intruder returning to maintain direct eye contact with the subject.

Behavioral responses of the monkeys were recorded using a head-mounted camera worn by the intruder and stationary cameras placed in the experimental room. The recorded behaviors were annotated using CAPTIV-L2100 software (TEA, France), subsequently classified by referencing the ethograms described in Izquierdo et al. (*64*) and Raper et al. (*65*) with some modification and were adapted to fit the specific behaviors observed in our subjects (**Table S2**). Behaviors were categorized into external (anxiety, excitation, defense, submission, affiliative) and internal (stereotyped behavior, exploratory behavior, others). To minimize observer bias, video recordings were coded independently by an expert in primate behavior and a naive non-expert observer. The mean of their scores was used for subsequent analyses.

### Standardized score calculation

To enable a comparative analysis of monkey behaviors, we calculated a standardized score for each behavioral parameter. This process involved two main steps: normalization of individual data points and subsequent calculation of the standardized score. First, raw behavioral data for each monkey from both the experimental and control group were normalized relative to the mean and standard error of the mean (SEM) of the control WT group’s data for each specific parameter:

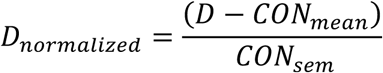

CON*_mean_* stands for the mean value of one parameter from the WT group, CON*_sem_* stands for the SEM of this parameter, “D” represents the specific per-monkey behavioral statistic in each parameter. We use two distinct normalized behavioral measures, the duration of time engaged in a specific category and the frequency of engagement in that category, to compute the final standardized score which was calculated as the sum of these two normalized values.

### Brain imaging analysis

#### MRI preparation

A total of 11 cynomolgus monkeys, comprising 6 WT individuals weighing 2.19±0.25 kg and 5 DISC1-LI individuals weighing 2.28 ± 0.29 kg, aged 2-2.5 years, underwent MRI scans. All subjects were housed individually in a light- and temperature-controlled colony room with feeding as described above and *ad libitum* access to water. None of the subjects had any prior implantation or cranial surgery.

On the day of the MRI scan, animals were food restricted until experiments were completed. Anesthesia was initiated by intramuscular injection of Alfaxalone (3 mg/kg) and Dexdomitor (dexmedetomidine hydrochloride; 5 µg/kg). Following sedation, the animals were transferred to the operating room in a secure transport cage. Endotracheal intubation was performed, and intravenous access was established to enable emergency intervention if needed. Once vital signs stabilized, the animals were transferred to the MRI suite for imaging. To investigate whether the alterations in brain functional connectivity were attributable to anesthesia or genetic mutation, an additional scan was conducted using Ketamine (5 mg/kg) in combination with Dexdomitor (5 µg/kg) and Atropine (0.05 mg/kg) to induce anesthesia. The interval between the two scans was at least one week.

#### MRI data acquisition

Both structural and functional MRI data were acquired using a whole-body 3T MRI scanner (Magnetom Verio; Siemens Healthineers AG, Erlangen, Germany). A custom-designed 8-channel cap-type radiofrequency coil, optimized for monkey anatomical skull structure obtained by CT, was used to enhance signal reception (Takashima Seisakusho Co., Ltd, Japan;). The animals were positioned in a prone posture without head fixation inside the scanner. General anesthesia was maintained throughout the procedure using isoflurane gas (1.0-1.5%) delivered in a mixture of air and oxygen. During the resting-state functional MRI (rsfMRI), the isoflurane level was lowered to 0.5%-0.6% to minimize its impact on spontaneous neural activity. Throughout the scanning session, physiological parameters, including heart rate, oxygen saturation (SpO_2_), respiratory rate, and end-tidal CO_2_ levels, were continuously monitored. The body temperature was maintained using thermal blankets and chemical heating pads to prevent hypothermia.

A 3D T1-weighted fast gradient echo imaging was conducted with the following parameters: TR = 11.8 ms, TE = 5.24 ms, flip angle = 20°, slices = 144, matrix = 192 × 192; FOV = 102 × 102 mm^2^; acquisition voxel size = 0.53 × 0.53 × 0.50 mm^3^. To facilitate effective correction of image distortions during post-processing, rsfMRI data were collected in both anterior-to-posterior (AP) and posterior-to-anterior (PA) phase encoding directions. A total of 1508 continuous 2D multiband echo-planar imaging (EPI) functional volumes were acquired, except for one WT subject for whom 1300 volumes were acquired, with the following parameters: TR = 1500 ms; TE = 28.4 ms; flip angle = 70°, slices = 44, matrix = 80 × 80; FOV = 96 × 96 mm^2^; acquisition voxel size = 1.2 × 1.2 × 1.2 mm^3^ with parallel imaging factor = 2 and multiband factor = 2 (*66*).

#### Image preprocessing for rsfMRI data

Preprocessing of the rsfMRI data was carried out integrating MATLAB with SPM12 (https://www.fil.ion.ucl.ac.uk/spm/software/spm12/) and FMRIB Software Library (FSL; https://fsl.fmrib.ox.ac.uk/fsl/docs/#/). After the correction for susceptibility-induced distortions and brain extraction by using TOPUP function of FSL, the brain mask was extracted by BET function. The distortion-corrected functional images were then spatially normalized to the ONPRC18 monkey template for the next analyses (*67*).

#### Independent component analysis (ICA)

Following spatial normalization, ICA was conducted for denoising purposes using FSL’s MELODIC toolbox. The initial 10 volumes of each dataset (AP and PA directions) were discarded to allow for signal stabilization, after which ICA was applied to the combined datasets. Subsequently, the filtered functional data resulting from group-level ICA were extracted for the accurate calculation of functional connectivity matrices.

#### Functional connectivity (FC) calculation

Following ICA, the filtered functional data from both AP and PA directions were prepared for FC analysis. Mean functional images corresponding to both directions were included in this preprocessing step. The datasets from these two directions were merged, and the T1-weighted brain image was aligned and normalized using SPM, ensuring proper anatomical correspondence between the functional and anatomical datasets. The functional data were subsequently normalized to the ONPRC 18 monkey template and atlas. Both AP and PA datasets were normalized with respect to the subject’s own T1-weighted image, maintaining spatial alignment across all subjects.

Prior to FC matrix calculation, the functional data were bandpass filtered to retain frequencies between 0.01 and 0.1 Hz, effectively removing high-frequency noise and low-frequency signal drift to focus on relevant resting-state fluctuations. Regions of interest (ROIs) were defined based on the ONPRC 18 atlas (Table S3), which was aligned with the normalized functional data. The time series were then extracted from each ROI, and pairwise Pearson’s correlations were calculated between all ROI pairs, generating a 55-by-55 FC matrix for each subject and the results were used for further analysis.

#### Graph theory analysis

A sparsity threshold of 0.05-0.30 was applied to FC matrices to binarize the connections, where correlations above the threshold were set to 1 (indicating the presence of a connection), and those below were set to 0 (indicating no connection). Self-connections (diagonal elements) were removed. Negative connections were treated as absent and hence not included in the binary matrices. The clustering coefficient, characteristic path length, small-worldness, global efficiency and local efficiency were calculated to compare the network properties between WT and DISC1-LI groups. To assess which brain regions function as hub-nodes in monkeys, eigenvector centrality was introduced. All properties were calculated using the NetworkX library in Python 3.13.

##### Clustering coefficient

The clustering coefficient *C_p_* of a network is a measure of the overall tendency of nodes to form tightly connected groups. It represents the average of the clustering coefficients of all individual nodes in the network. It is calculated as:

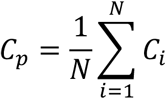

where *C_i_* is the clustering coefficient of node *i*, as given by:

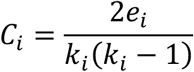

Here, *e_i_* is the number of edges between the neighbors of node *i*, and *k_i_* is the degree of node *i*.

##### Characteristic path length

The characteristic path length *L* is the average shortest path between all pairs of nodes in a network, defined as:

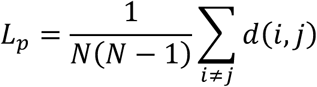

where *d*(*i*, *j*) is the shortest path between nodes *i* and *j*, and *N* is the total number of nodes in the network.

##### Small-worldness

Small-worldness of brain networks is often characterized by high clustering and short path lengths, a balance that allows for both segregated and integrated processing of information (*68*). To assess the small-world characteristics of the monkey brain networks, we calculated the small-world coefficient *σ*, defined as:

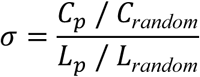

where *C_p_* and *L_p_* are the clustering coefficient and characteristic path length of the real network, respectively, and *C_random_*and *L_random_*are the corresponding values for a random network of the same size and degree distribution. The networks *C_p_* and *L_p_* were compared with 100 matched random networks (each with the same number of nodes and edges and degree distribution), resulting in the normalized clustering coefficient *γ* = *C_p_* / *C_random_*and normalized characteristic path length *λ* = *L_p_* /*L_random_*. A network is considered small world if *σ*>1. Random networks were generated by randomly rewiring the edges of the real networks while preserving the degree distribution.

##### Global efficiency

Global efficiency is a measure of how efficient information can be exchanged across the entire network. It is calculated as the average length of the inverse shortest path lengths between all pairs of nodes. It reflects the overall ability of the network to transfer information globally. The definition of global efficiency is:

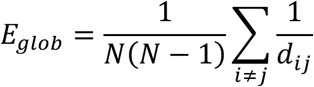

Where *N* is the total number of nodes in the network. *d_ij_* is the shortest path length between node *i* and *j*. If *d_ij_* is infinite (nodes are not connected), 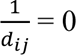.

##### Local efficiency

Local efficiency measures how well information is exchanged within the local neighborhood of a node. It reflects the efficiency of the subgraph formed by the immediate neighbors of a node when that node is removed. It is defined as the average global efficiency of the subgraphs of neighbors. The local efficiency is defined as:

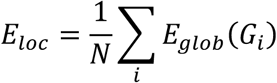

where *G_i_* is the subgraph consisting of the neighbors of node *i*. *E*_g*lob*_(*G_i_*) is the global efficiency of the subgraph *G_i_*.

##### Eigenvector centrality

Eigenvector centrality is an indictor to measure the importance of a node in a network by considering not only the number of its connections but also the importance of the nodes it is connected to. Mathematically, the centrality *x* of node *i* is defined as:

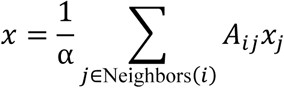

where *A* is the adjacency matrix, *α* is the largest eigenvalue, and *x* is the eigenvector corresponding to *α*. Nodes connected to highly central nodes will themselves have higher centrality scores.

### Statistical analyses

Statistical comparisons between groups were conducted using GraphPad Prism 10 (GraphPad Software, Boston, MA, USA) or Python (version 3.13). The specific statistical tests used for each analysis are described in the respective figure legends. Statistical significance was defined as *P* < 0.05.

## Acknowledgments

We thank Yushi Kitada, Kaoru Isa, Sayaka Matsushita, Yuto Fukushima and Masashi Nakamura for their animal care and experimental assistance. We also thank Dr. Melissa Salgado for her critical reading and English revision of the manuscript.

## Data and Materials Availability

All data needed to evaluate the conclusions in the paper are present in the paper and/or the Supplementary Materials.

