## Supplementary information for "Adolescent prodromal phenotypes of psychiatric disorders: insights from a novel DISC1-perturbed monkey model"

Lan *et al*.

*Corresponding author:
Akira Sawa
Tadashi Isa

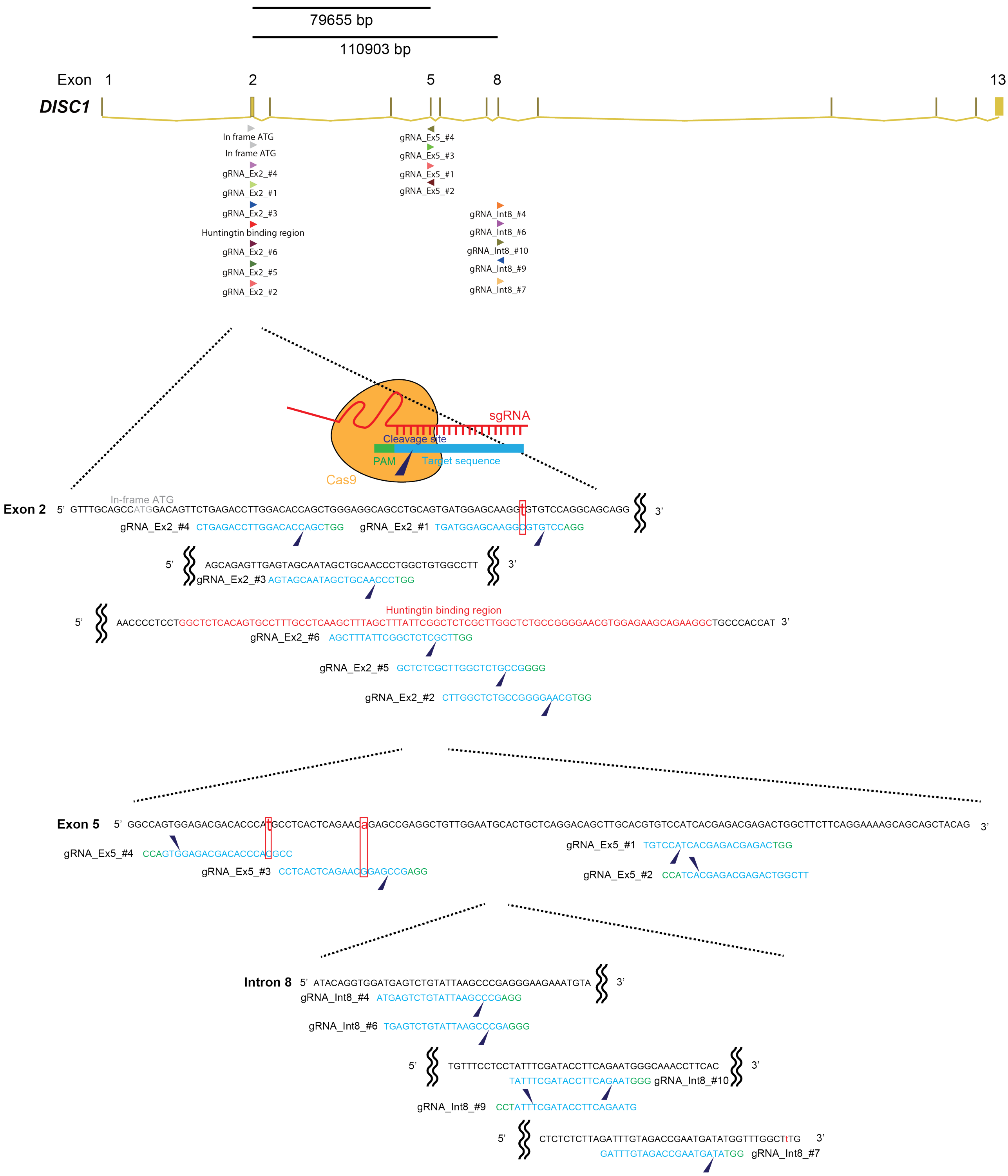

**Fig. S1. Design of sgRNAs for *DISC1* manipulation.** Schematic representation of the macaque *DISC1* locus and the locations of the designed guide RNAs targeting Exon 2, Exon 5, and Intron 8. Lowercase letters in the nucleotide sequences indicate genetic polymorphisms.

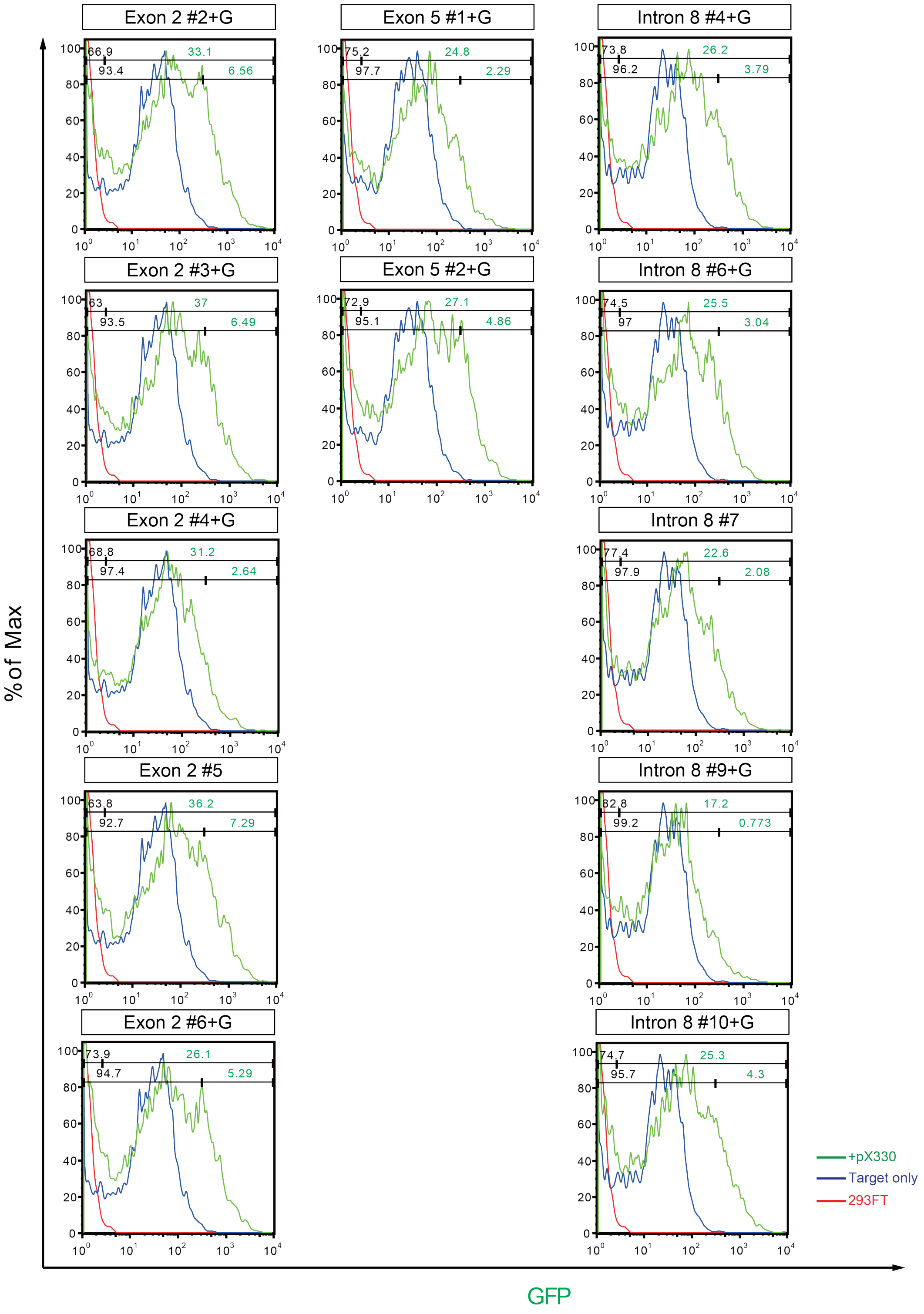

**Fig. S2. Single-strand annealing (SSA) assay.** EGFP signals 2 days after 293FT cells were co-transfected with an EGxxFP vector and CRISPR/Cas9 vectors. The numbers in the histograms indicate the percentages of cells exhibiting weak and strong GFP-positive signals under the +pX330 condition.

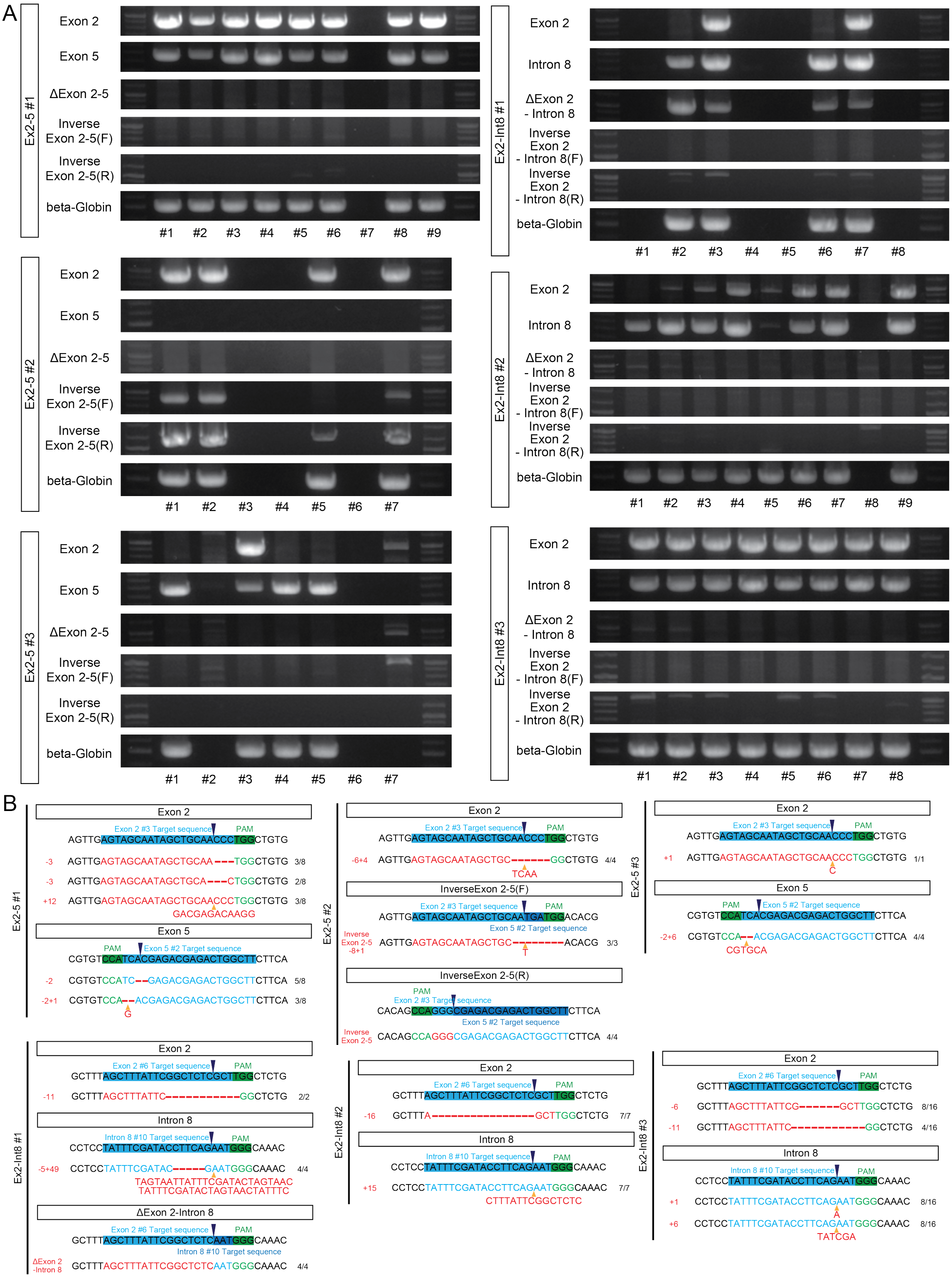

**Fig. S3. Genotyping the DISC1-LI embryos.** (**A**) Genomic PCR for genotyping DISC1-LI embryos. The numbers in the horizontal axis of each panel indicate individual blastomeres sampled from each embryo. (**B**) Sequences of sgRNA targets in the genomes of the DISC1-LI embryos.

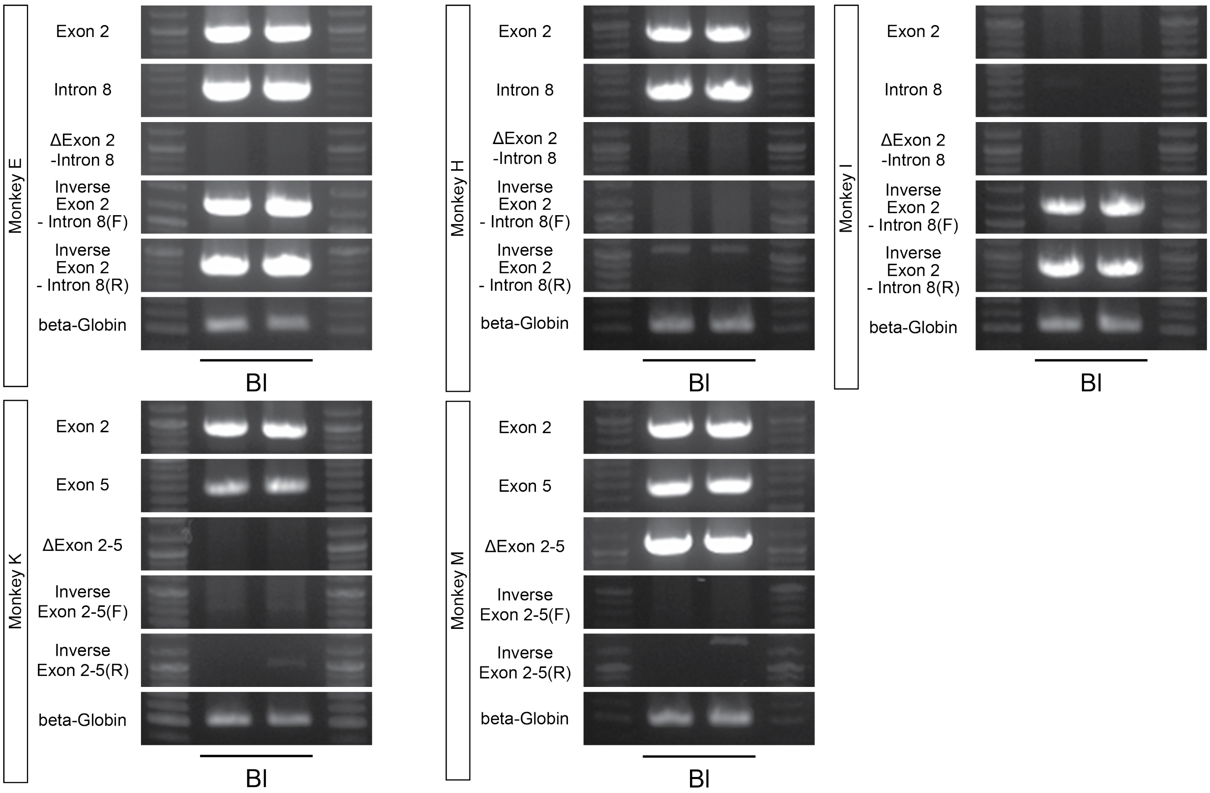

**Fig. S4. Genotyping the DISC1-LI monkeys.** Genomic PCR for genotyping DISC1-LI monkeys.

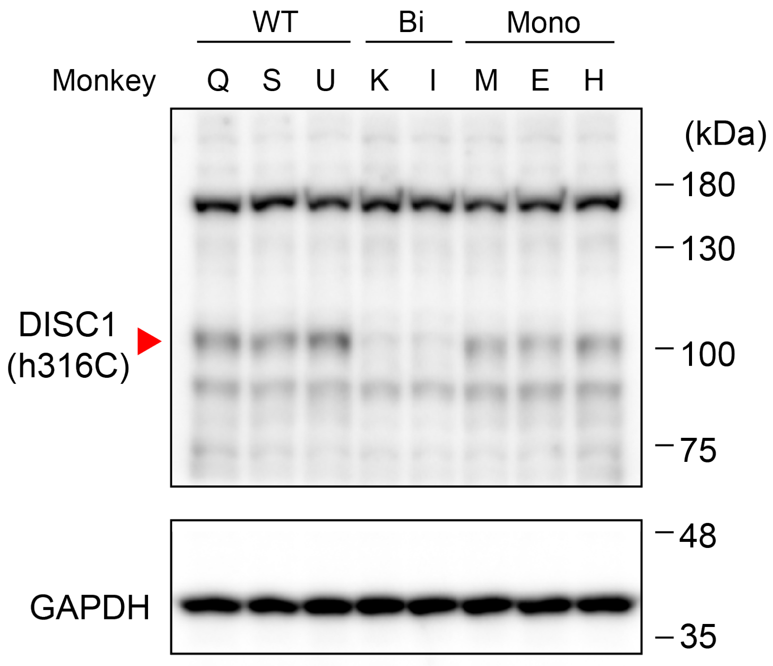

**Fig. S5. Western blotting for DISC1 in iPS cells derived from monkeys**. GAPDH was a loading control.

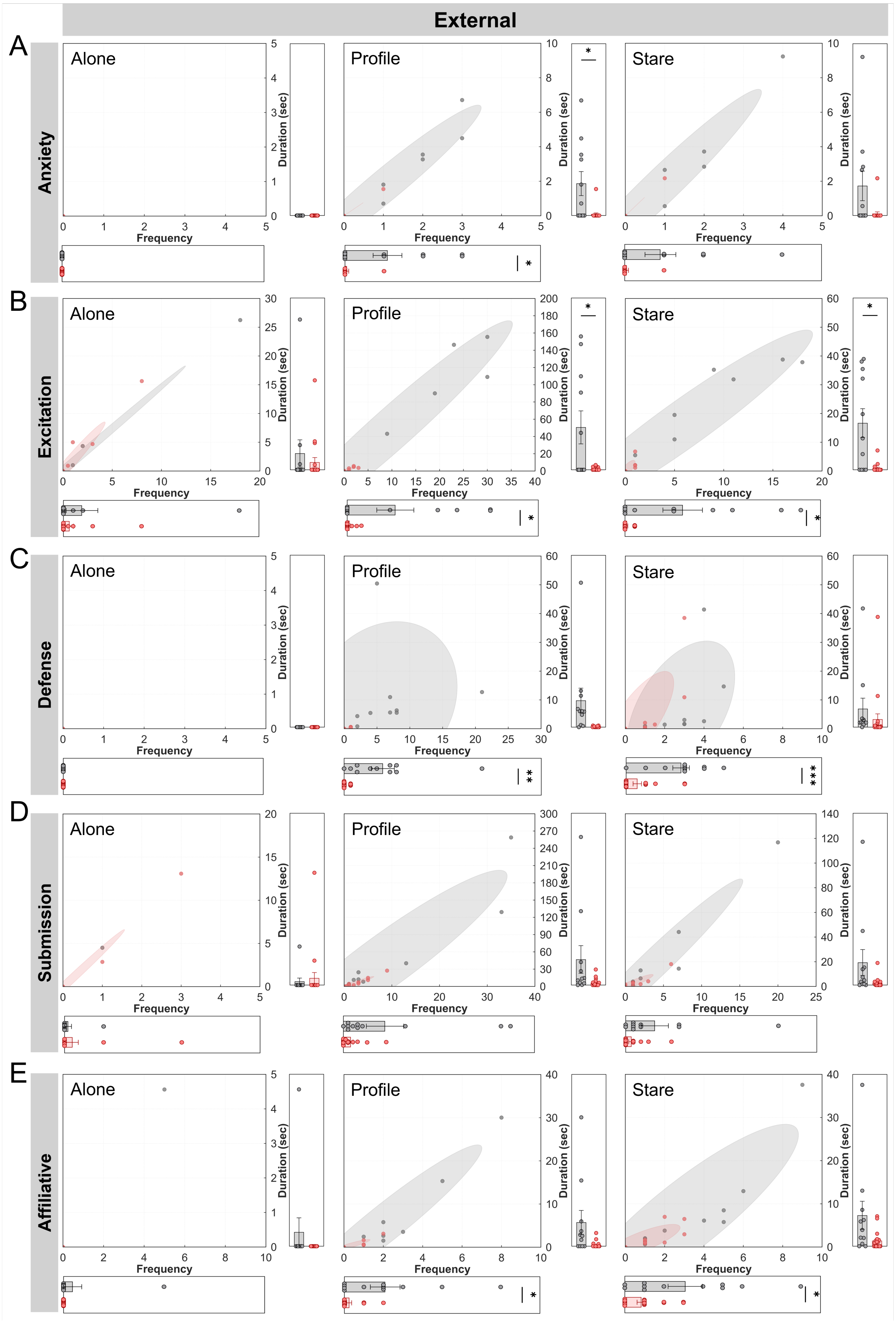
External
A
Anxiety
Alone
Profile
Stare
B
Excitation
C
Defense
D
Submission
E
Affiliative
Frequency
Duration (sec)

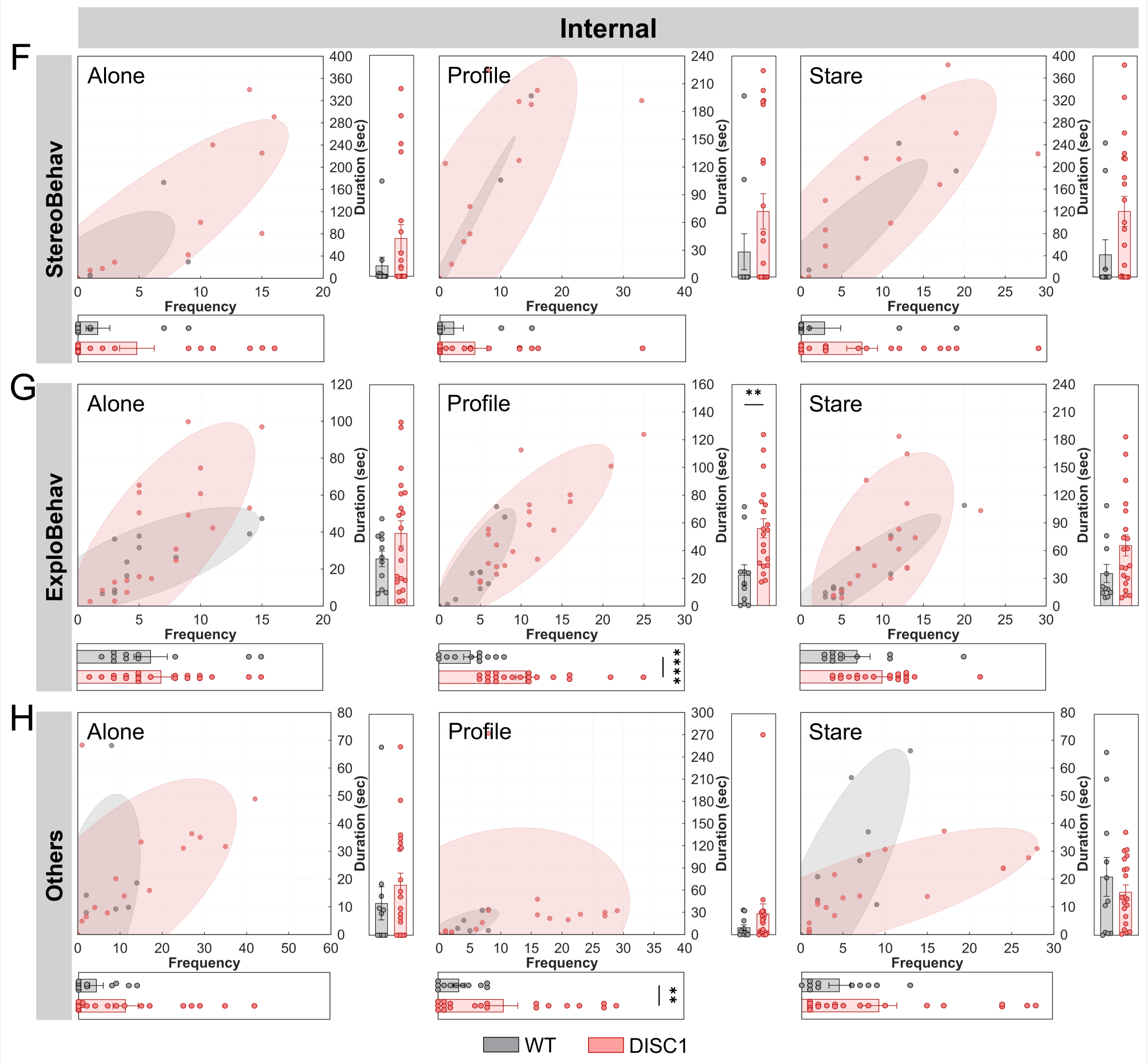

**Fig. S6. Detailed behavioral activity of monkeys during the HIT.** (**A**) Anxiety-related behavior (*Profile*: duration, $*P = 0.0289$; frequency, $*P = 0.0181$). (**B**) Excitation-related behavior (*Profile*: duration, $*P = 0.0299$; frequency, $*P = 0.0321$; *Stare*: duration, $*P = 0.0108$; frequency, $*P = 0.018$). (**C**) Defense-related behavior (*Profile*: frequency, $**P = 0.0087$; *Stare*: frequency, $***P = 0.0003$). (**D**) Submission-related behavior. n.s. (**E**) Affiliative behavior (*Profile*: frequency, $*P = 0.036$; *Stare*: frequency, $*P = 0.0304$). (**F**) Stereotyped behavior. n.s. (**G**) Exploratory behavior (*Profile*: duration, $**P = 0.0027$; frequency, $****P < 0.0001$). (**H**) Other behaviors (*Profile*: frequency, $**P = 0.0085$). Unpaired t-test with Welch's correction.

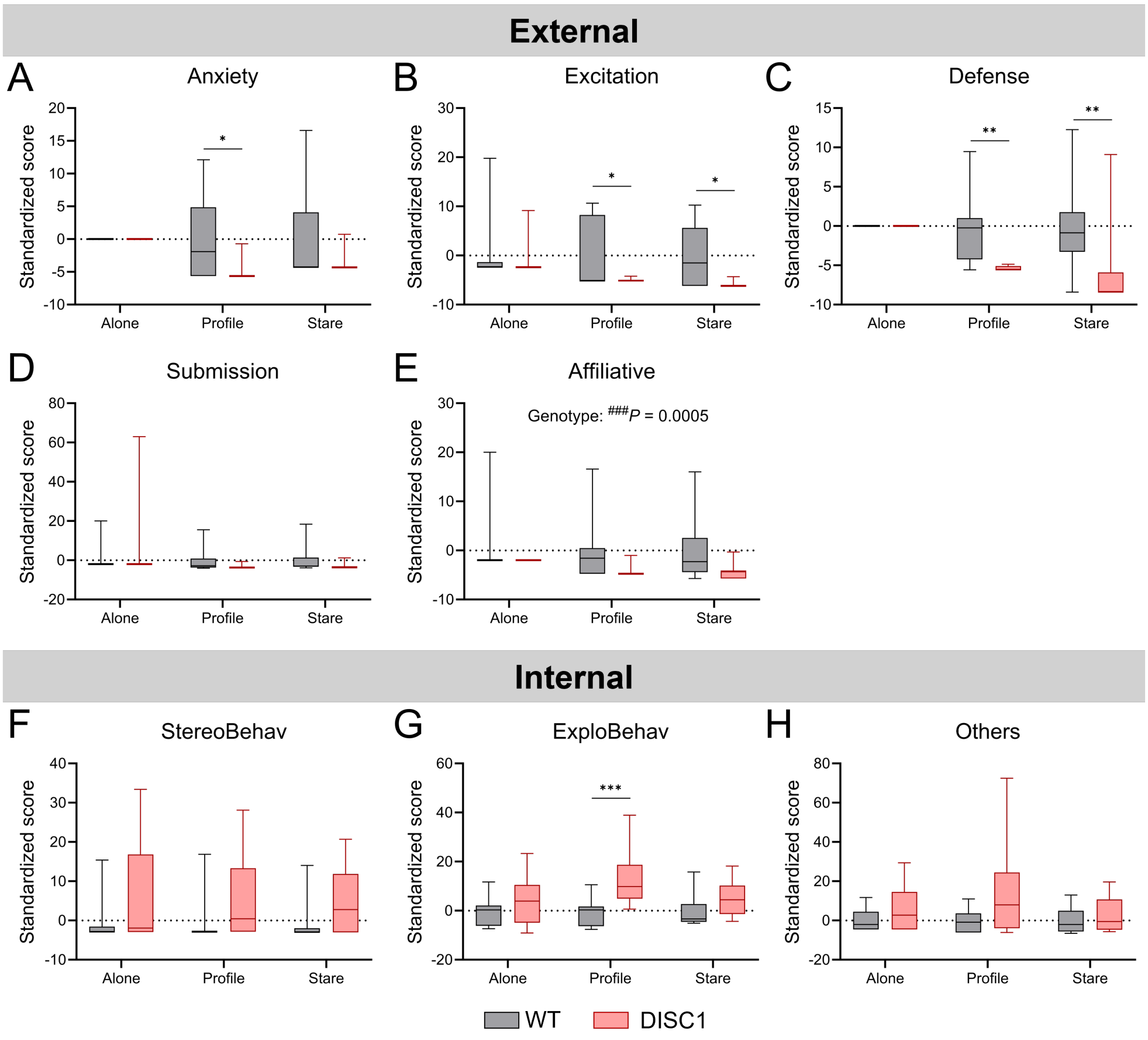

**Fig. S7. Standardized score of all behavioral categories in monkeys during the HIT.** (**A**) Anxiety-related behavior. *Profile*, $^{*}P = 0.022$ (Genotype × Phase, $^{\#\#}P = 0.006$; Genotype, $^{\#\#\#}P = 0.0002$; Phase, $^{\#\#}P = 0.0093$). (**B**) Excitation-related behavior. *Profile*, $^{*}P = 0.0299$; *Stare*, $^{*}P = 0.0131$ (Genotype × Phase, $^{\#}P = 0.0115$; Genotype, $^{\#\#}P = 0.0025$; Phase, $^{\#}P = 0.0322$). (**C**) Defense-related behavior. *Profile*, $^{**}P = 0.0057$; *Stare*, $^{**}P = 0.0059$ (Genotype × Phase, $^{\#\#\#}P = 0.0005$; Genotype, $^{\#\#}P < 0.0001$; Phase, $^{\#\#}P = 0.0014$). (**D**) Submission-related behavior. n. s. (**E**) Affiliative behavior. Genotype, $^{\#\#\#}P = 0.0005$. (**F**) Stereotyped behavior. n. s. (**G**) Exploratory behavior. *Profile*, $^{***}P = 0.0002$ (Genotype × Phase, $^{\#}P = 0.0407$; Genotype, $^{\#\#}P = 0.0031$; Phase, $^{\#}P = 0.0481$). (H) Other behaviors. n. s. Two-way RM ANOVA with Greenhouse-Geisser correction, followed by Tukey's HSD test.

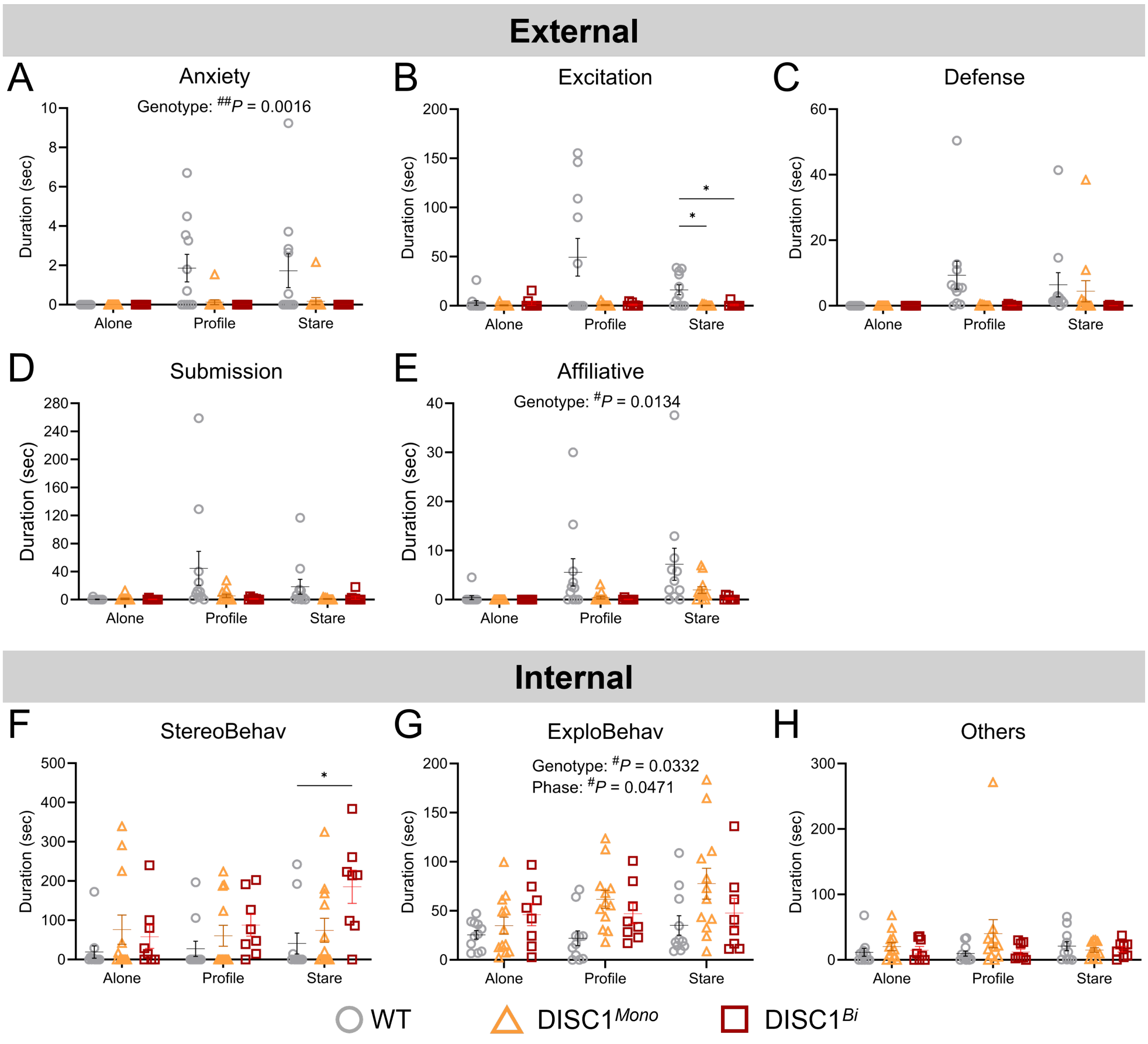

**Fig. S8**. **Behavioral duration of the HIT in all genotypes.** WT, $N = 4$ (total trials = 11); DISC1$^{Mono}$, $N = 3$ (total trials = 12); DISC1$^{Bi}$, $N = 2$ (total trials = 8). (**A**) Anxiety-related behavior. Genotype, $^{\#\#}P = 0.0016$ (WT vs. DISC1$^{Mono}$, $^{\#\#}P = 0.0044$; WT vs. DISC1$^{Bi}$, $^{\#\#}P = 0.0051$). (**B**) Excitation-related behavior. Stare: WT vs. DISC1$^{Mono}$, $^{*}P = 0.0252$; WT vs. DISC1$^{Bi}$, $^{*}P = 0.0293$ (Genotype × Phase, $^{\#\#\#}P = 0.0008$; Genotype, $^{\#\#}P = 0.006$; Phase, $^{\#\#}P = 0.0314$). (**C**) Defense-related behavior. n. s. (**D**) Submission-related behavior. Genotype × Phase, $^{\#}P = 0.0443$. (**E**) Affiliative behavior. Genotype, $^{\#}P = 0.0134$ (WT vs. DISC1$^{Mono}$, $^{\#}P = 0.0364$; WT vs. DISC1$^{Bi}$, $^{\#}P = 0.0231$). (**F**) DISC1$^{Bi}$ showed more stereotyped behavior compared to WT ($^{*}P = 0.0328$). Genotype × Phase, $^{\#\#\#}P = 0.0003$; Phase, $^{\#\#}P = 0.0002$. (**G**) Exploratory behavior. Genotype, $^{\#}P = 0.0332$ (WT vs. DISC1$^{Mono}$, $^{\#}P = 0.0266$); Phase ($^{\#}P = 0.0471$). (H) Other behaviors. n. s. Two-way RM ANOVA with Greenhouse-Geisser correction, followed by Tukey's HSD test.

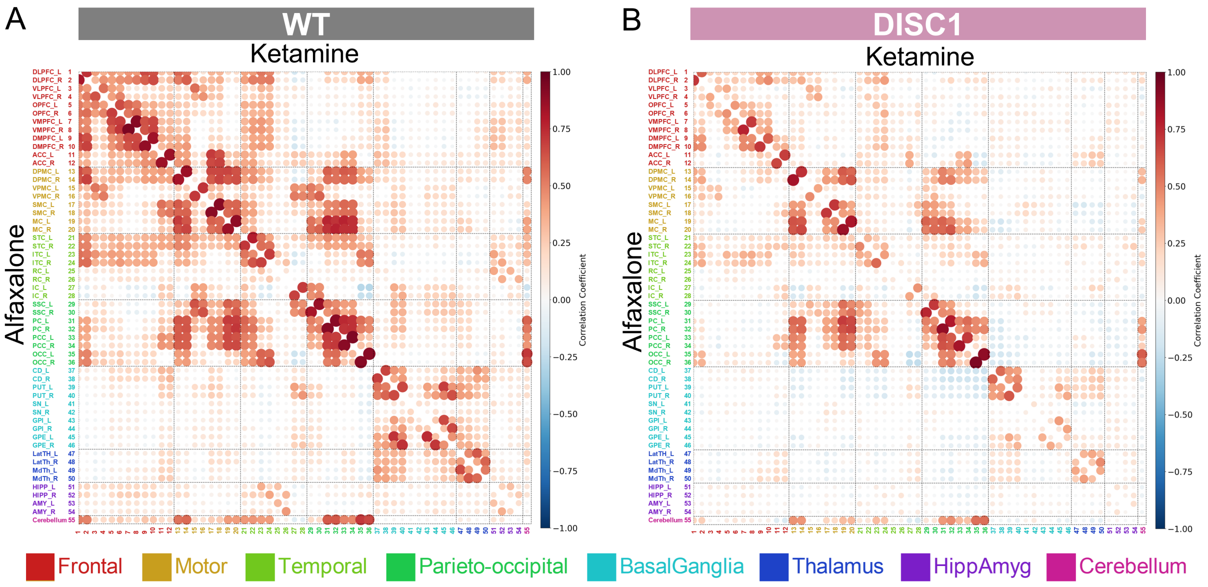

**Fig. S9. Comparison of FC under alfaxalone and ketamine anesthesia.** (**A**) FC matrix for WT under alfaxalone ($N = 6$) and ketamine anesthesia ($N = 4$). (**B**) FC matrix for DISC1-LI monkeys under alfaxalone and ketamine anesthesia ($N = 5$).

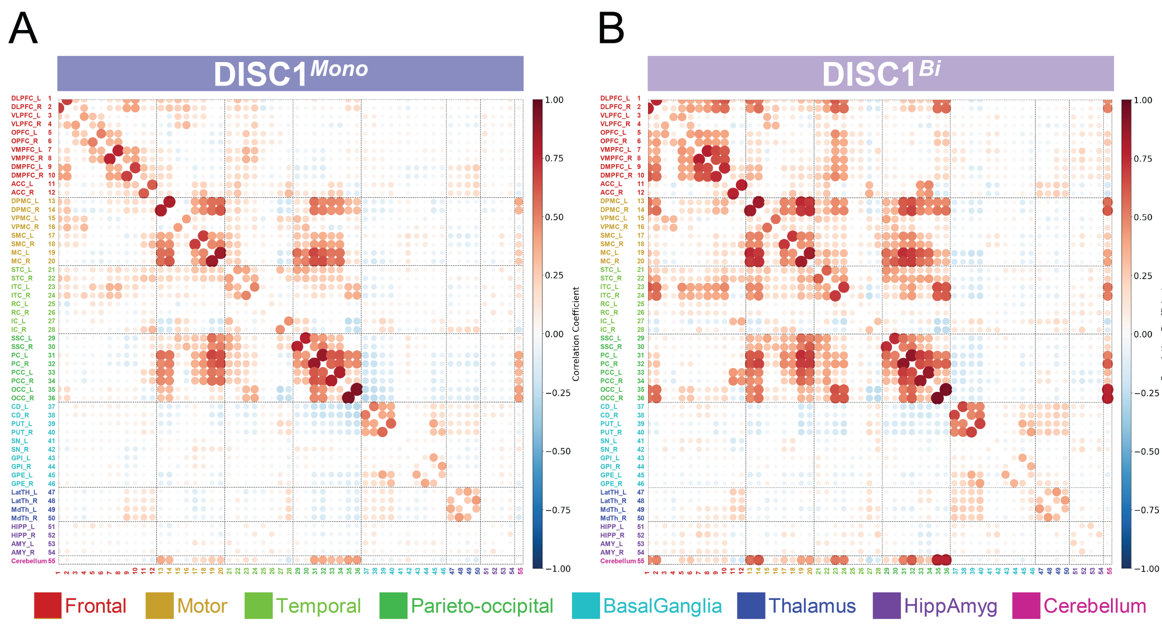

**Fig. S10. FC in different mutation types.** (**A**) FC matrix for *DISC1* monoallelic mutation (*N* = 3). **(B)** FC matrix for *DISC1* biallelic mutation (*N* = 2). Both mutations led to great reductions in global functional connectivity.

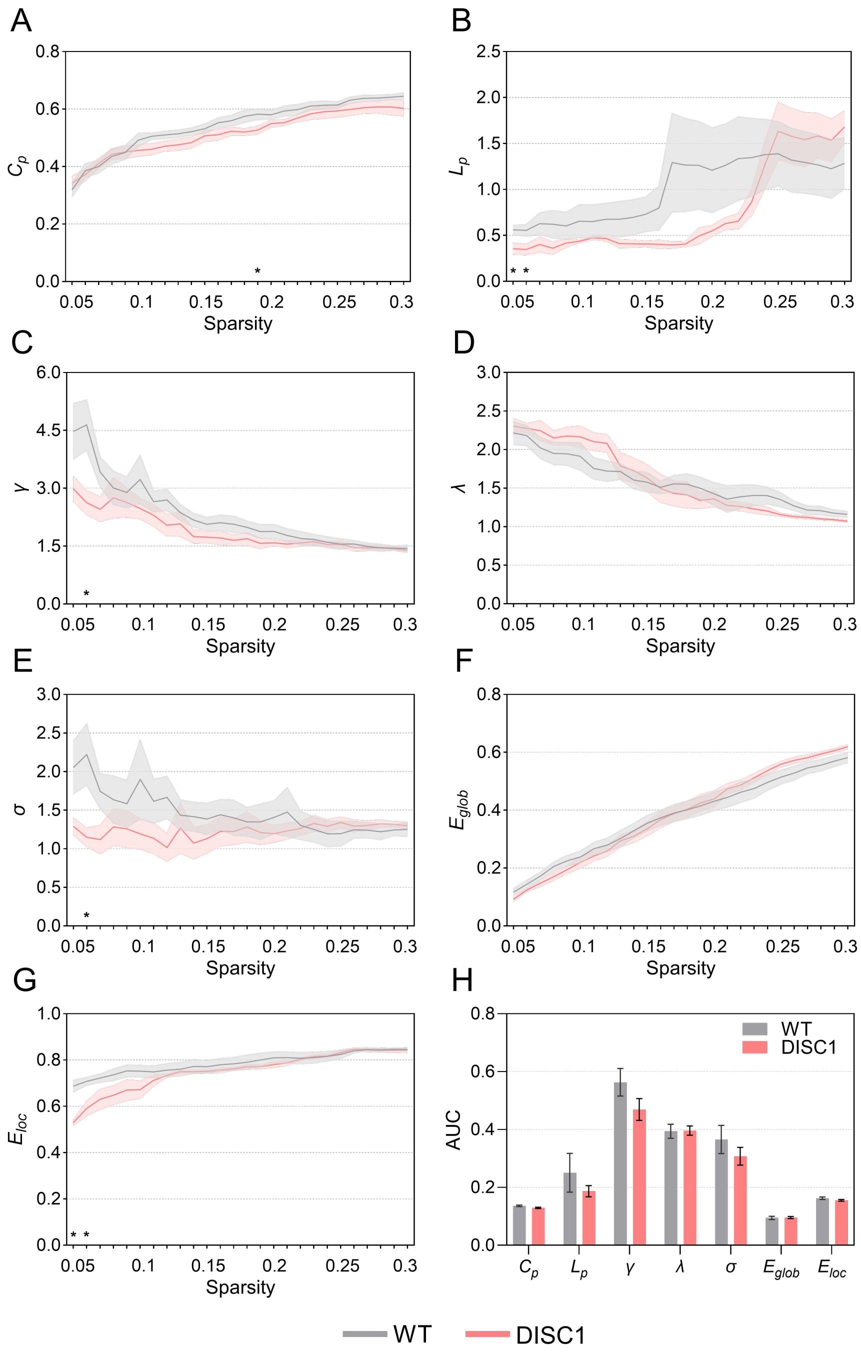

**Fig. S11. Small-world properties of functional brain networks in WT and DISC1-LI monkeys.** (**A** to **G**) Small-world properties across sparsity thresholds (0.05 ≤ S ≤ 0.30). (**A**) Clustering coefficient. (**B**) Characteristic path length. (**C**) Normalized clustering coefficient. (**D**) Normalized characteristic path length. (**E**) Small-worldness. (**F**) Global efficiency. (**G**) Local efficiency. (**H**) Area under the curves. Significant differences are indicated as $*P < 0.05$. Error bars represent standard errors of the mean.

**Table S1. Oligonucleotides used in this study.**

| Names | Forward | Reverse | For |
|---|---|---|---|
| DISC1_Ex2_ #2+G_F,R | CACCGCTTGGCTCTGCC GGGGAACG | AAACCGTTCCCCGGCAG AGCCAAGC | sgRNA cloning (Ex2) |
| DISC1_Ex2_ #3+G_F,R | CACCGAGTAGCAATAGC TGCAACCC | AAACGGGTTGCAGCTAT TGCTACTC | sgRNA cloning (Ex2) |
| DISC1_Ex2_ #4+G_F,R | CACCGCTGAGACCTTGG ACACCAGC | AAACGCTGGTGTCCAAG GTCTCAGC | sgRNA cloning (Ex2) |
| DISC1_Ex2_ #5_F,R | CACCGCTCTCGCTTGGCT CTGCCG | AAACCGGCAGAGCCAAG CGAGAGC | sgRNA cloning (Ex2) |
| DISC1_Ex2_ #6+G_F,R | CACCGAGCTTTATTCGG CTCTCGCT | AAACAGCGAGAGCCGA ATAAAGCTC | sgRNA cloning (Ex2) |
| DISC1_Ex5_ #1+G_F,R | CACCGTGTCCATCACGA GACGAGAC | AAACGTCTCGTCTCGTG ATGGACAC | sgRNA cloning (Ex5) |
| DISC1_Ex5_ #2+G_F,R | CACCGAAGCCAGTCTCG TCTCGTGA | AAACTCACGAGACGAGA CTGGCTTC | sgRNA cloning (Ex5) |
| DISC1_Int8_ #4+G_F,R | CACCGATGAGTCTGTAT TAAGCCCG | AAACCGGGCTTAATACA GACTCATC | sgRNA cloning (Int8) |
| DISC1_Int8_ #6+G_F,R | CACCGTGAGTCTGTATT AAGCCCGA | AAACTCGGGCTTAATAC AGACTCAC | sgRNA cloning (Int8) |
| DISC1_Int8_ #7_F,R | CACCGATTTGTAGACCG AATGATA | AAACTATCATTCGGTCT ACAAATC | sgRNA cloning (Int8) |
| DISC1_Int8_ #9+G_F,R | CACCGCATTCTGAAGGT ATCGAAAT | AAACATTTCGATACCTT CAGAATGC | sgRNA cloning (Int8) |
| DISC1_Int8_ #10+G_F,R | CACCGTATTTCGATACCT TCAGAAT | AAACATTCTGAAGGTAT CGAAATAC | sgRNA cloning (Int8) |
| DISC1_Ex2_t arget_EcoRI_ F, NheI_R | GAATTCCCAGAGAGTGT GGCCTTGACCTGAGAG | GCTAGCGTCCTCTGAGA CCTGTGTAGCCAAGAG | SSA assay (Ex2) |

| | | | |
|---|---|---|---|
| DISC1_Ex5_target_EcoRI_F, NheI_R | GAATTCCAGCTAAGTACCCTCCTAAGTATGAGAACAG | GCTAGCGCTACACTCAAGGAAAGTAAGACCCAC | SSA assay (Ex5) |
| DISC1_Int8_target_EcoRI_F, NheI_R | GAATTCATTGAACACAGTCTTGCTTCTGAGTTAGG | GCTAGCGAAACCACCCTCATGATCCAAACACCTC | SSA assay (Int8) |
| Cyn_beta-globin_genome_F, R | AAGGTGAACGTGGATGAAGTTGGTGG | TGTCTCTTCCCCATTCTAAACTGTACCCTG | Genotyping (beta-globin) |
| DISC1_Ex2_geno_F, target_NheI_R | GGGATTTTCATAACGGATTGGCCCCAG | GCTAGCGTCCTCTGAGACCTGTGTAGCCAAGAG | Genotyping (Ex2) |
| DISC1_Ex5_target_EcoRI_F, NheI_R | GAATTCCAGCTAAGTACCCTCCTAAGTATGAGAACAG | GCTAGCGCTACACTCAAGGAAAGTAAGACCCAC | Genotyping (Ex5) |
| DISC1_Int8_target_EcoRI_F, NheI_R | GAATTCATTGAACACAGTCTTGCTTCTGAGTTAGG | GCTAGCGAAACCACCCTCATGATCCAAACACCTC | Genotyping (Int8) |
| DISC1_Ex2_geno_F, Ex5_target_NheI_R | GGGATTTTCATAACGGATTGGCCCCAG | GCTAGCGCTACACTCAAGGAAAGTAAGACCCAC | Genotyping (delta Ex2-5) |
| DISC1_Ex2_geno_F, Ex5_target_EcoRI_F | GGGATTTTCATAACGGATTGGCCCCAG | GAATTCCAGCTAAGTACCCTCCTAAGTATGAGAACAG | Genotyping (inversion Ex2-5 (F)) |
| DISC1_Ex2_geno_R, Ex5_target_NheI_R | GTGACAGAGAAGACAAAAGGGAGGGTAGATC | GCTAGCGCTACACTCAAGGAAAGTAAGACCCAC | Genotyping (inversion Ex2-5 (R)) |
| DISC1_Ex2_geno_F, | GGGATTTTCATAACGGATTGGCCCCAG | GCTAGCGAAACCACCCTCATGATCCAAACACCTC | Genotyping (delta Ex2-Int8) |

| | | | |
|---|---|---|---|
| Int8_target_NheI_R | | | |
| DISC1_Ex2_geno_F, Int8_target_EcoRI_F | GGGATTTTCATAACGGATTGGCCCCAG | GAATTCATTGAACACAGTCTTGCTTCTGAGTTAGG | Genotyping (inversion Ex2-Int8 (F)) |
| DISC1_Ex2_geno_R, Int8_target_NheI_R | GTGACAGAGAAGACAAAAGGGAGGGTAGATC | GCTAGCGAAACCACCCTCATGATCCAAACACCTC | Genotyping (inversion Ex2-Int8 (R)) |

**Table S2. Behavioral ethogram for human intruder test**

| | Behavior categories | Description |
|---|---|---|
| External | **(1) Anxiety** | |
| | Body shake | Shaking the whole body, or the head and shoulders. |
| | Teeth grind | Repetitive, audible rubbing of upper and lower teeth. |
| | Yawn | Opening mouth widely, exposing teeth. |
| | **(2) Excitation** | |
| | Cage aggression | Hitting or shaking the cage, slamming the body against it. |
| | Threat | Opening the mouth without showing teeth, quick head movements, moving eyebrows. |
| | Self-biting | Biting one's own arms or legs. |
| | **(3) Defense** | |
| | Startle | Sudden, quick movements; moving away from a stimulus. |
| | Eye/ Head aversion | Avoiding eye contact, turning away the gaze or head. |
| | **(4) Submission** | |
| | Lip smack | Protruding lips and moving them rapidly. |
| | Grimace | Pulling back the lips to expose the teeth. |
| | Present | Raising the tail to expose the rear. |
| | **(5) Affiliative** | |
| | Eye contact | Making eye contact. |
| | Move toward | Approaching the stimulus. |
| | Touch | Touching with hands or feet. |
| | Social grooming | Grooming a person's hand or clothing. |
| | Soft mouth | Gently biting. |
| | Ear flapping | Flapping ears. |
| | Mounting | Pressing the genitals or mounting on a hand. |
| Internal | **(6) Stereotyped behavior** | |
| | Pacing | Walking back and forth along the same path rhythmically. |
| | Motor stereotype | Rocking the body back and forth, spinning in circles, or performing flips. |
| | **(7) Exploratory behavior** | |
| | Manual/ Oral exploration | Touching various objects, licking, putting things in the mouth, or exploring. |
| | **(8) Others (not directed towards the stimulus)** | |
| | Self-directed activities | Scratching the body with hands or feet, self-grooming. |
| | Vocalization | |
| | ・coo call | A soft, clear, low-pitched sound, “oooh”. |
| | ・scream | A loud, high-pitched sound or chirp, “screech”. |

**Table S3. Region of interest for rsfMRI**

| Label # | Atlas Region | Abbreviation | Consolidated Atlas Regions |
|---|---|---|---|
| 1, 2 | Dorsolateral Prefrontal Cortex | DLPFC_L/R | Frontal |
| 3, 4 | Ventrolateral Prefrontal Cortex | VLPFC_L/R | |
| 5, 6 | Orbitofrontal Prefrontal Cortex | OPFC_L/R | |
| 7, 8 | Ventromedial Prefrontal Cortex | VMPFC_L/R | |
| 9, 10 | Dorsomedial Prefrontal Cortex | DMPFC_L/R | |
| 11, 12 | Anterior Cingulate Cortex | ACC_L/R | |
| 13, 14 | Dorsal Premotor Cortex | DPMC_L/R | Motor |
| 15, 16 | Ventral Premotor Cortex | VPMC_L/R | |
| 17, 18 | Supplementary Motor Cortex | SMC_L/R | |
| 19, 20 | Primary Motor Cortex | MC_L/R | |
| 21, 22 | Superior Temporal Cortex | STC_L/R | Temporal |
| 23, 24 | Inferior Temporal Cortex | ITC_L/R | |
| 25, 26 | Rhinal Cortex | RC_L/R | |
| 27, 28 | Insular Cortex | IC_L/R | |
| 29, 30 | Somatosensory Cortex | SSC_L/R | Parieto-occipital |
| 31, 32 | Parietal Cortex | PC_L/R | |
| 33, 34 | Posterior Cingulate Cortex | PCC_L/R | |
| 35, 36 | Occipital Cortex | OCC_L/R | |
| 37, 38 | Caudate | CD_L/R | Basal Ganglia |
| 39, 40 | Putamen | PUT_L/R | |
| 41, 42 | Substantia nigra | SN_L/R | |
| 43, 44 | Globus Pallidus Internal | GPI_L/R | |
| 45, 46 | Globus Pallidus External | GPE_L/R | |
| 47, 48 | Lateral Thalamus | LatTh_L/R | Thalamus |
| 49, 50 | Medial Thalamus | MdTh_L/R | |
| 51, 52 | Hippocampus | HIPP_L/R | Hippocampus |
| 53, 54 | Amygdala | AMY_L/R | Amygdala |
| 55 | Cerebellum | - | Cerebellum |
